# Tumor proliferation and invasion are coupled through cell-extracellular matrix friction

**DOI:** 10.1101/2022.11.15.516548

**Authors:** Ashleigh J. Crawford, Clara Gomez-Cruz, Gabriella C. Russo, Wilson Huang, Isha Bhorkar, Arrate Muñoz-Barrutia, Denis Wirtz, Daniel Garcia-Gonzalez

## Abstract

Cell proliferation and invasion are two key drivers of tumor progression and are traditionally considered two independent cellular processes regulated by distinct pathways. Through *in vitro* and *in silico* methods, we provide evidence that these two processes are intrinsically coupled through matrix-adhesion friction. Using novel tumor spheroids, we show that both tumor cell proliferation and invasion are limited by a volumetric carrying capacity of the system, i.e. maximum spatial cell concentration supported by the system’s total cell count, nutrient consumption rate, and collagen gel mechanical properties. To manipulate these phenotypes in breast cancer cells, we modulate the expression of E-cadherin and its associated role in adhesion, invasion, and proliferation. We integrate these results into a mixed-constitutive formulation to computationally delineate the contributions of cellular and extracellular adhesion, stiffness, and mechanical properties of the extracellular matrix (ECM) to the proliferative and invasive fates of breast cancer tumor spheroids. Both approaches conclude that the dominant drivers of tumor fate are system properties modulating cell-ECM friction, such as E-cadherin dependent cell-ECM adhesion and matrix pore size.

## 1. Introduction

As the field of cancer research has rapidly advanced over the last fifty years, the hallmarks of cancer have been discussed, defined, and revisited. With each iteration, these hallmarks consistently focus on two fundamental cellular processes: proliferation and invasion/migration [1, 2]. Associated tumor growth and metastasis are regulated by these two processes. This statement is an oversimplification, however, as each process can be subdivided into an array of phenotypes contributing to tumor fate. Proliferation involves the balance of cell proliferation, cell death, and metabolic rewiring to sustain these fates; migration/invasion includes both single-cell and collective invasion into the tissue stroma, as well as directed cell migration. Nevertheless, according to the “go or grow” hypothesis, these events are traditionally considered mutually exclusive [3, 4, 5]. Some groups have confirmed this hypothesis by observing cancer cells that alternate between highly motile and highly proliferative phenotypes [6, 7, 8, 9], while others report a correlation between tumor proliferation and invasion [10, 11, 12].

E-cadherin (E-cad) has long been associated with the epithelial-to-mesenchymal transition (EMT), where the loss of E-cad expression is associated with a shift to a mesenchymal phenotype [13, 14]. E-cad also increases cell-cell adhesion via the formation of adherens junctions. The transition to a mesenchymal phenotype and decrease in cell-cell adhesion associated with the loss of E-cad expression encourage single-cell invasion, while E-cad expression favors collective cell invasion [15]. More recent studies on E-cad expression in invasive ductal carcinoma (IDC) have provided evidence that E-cad also alters the proliferative phenotype of breast cancer cells through EGFR/MEK activation [16]. E-cad positive (E-cad+) cells are hyper-proliferative compared to E-cad negative (E-cad-) cells. IDC patients with tumors that are E-cad+ have a lower survival rate than those with E-cad-tumors and suffer from metastatic disease more frequently [16, 17]. Thus, this paradoxical role of E-cad in the clinical setting has led to a re-examination of its classification and a focus on the interactions between the tumor and its microenvironment.

IDC tumors remodel the tumor microenvironment and deposit type I collagen [18]. This increase in collagen density changes the physical and mechanical properties of the surrounding stroma which in turn alters the cell-extracellular matrix (ECM) interactions. *In vitro* studies have demonstrated that fiber alignment, pore size, and cell speed all decrease with increasing collagen concentration [19]. However, the elastic modulus changes non-monotonically with collagen concentration, leading to irregularities that are difficult to predict [19]. Our overarching hypothesis is that proliferation and collective invasion are coupled and regulated by both cell-cell and cell-ECM interactions. The constraints enforced by the extracellular system ultimately determine the fate of the tumor. Thus, we observe a subdued proliferative response to increases in matrix stiffness in 3D culture. When high matrix density in the surrounding tumor microenvironment prevents cell invasion, proliferation is limited by the volumetric carrying capacity of the tumor.

Here, we develop and implement a computational model to unravel the intertwined dynamics of cell proliferation and invasion in an IDC tumor spheroid. We study cell proliferation and invasion as highly coordinated events: tumor proliferation can be limited by low invasion rates and encouraged by high invasion rates, and tumor invasion requires support from a high proliferation rate or will be limited by low cell counts. We use a novel oil-in-water droplet technology to suspend a dense cluster of E-cad-or E-cad+ MDA-MB-231 cells in matrices of increasing type I collagen density [20]. The implantation of the spheroids into matrices of excess volume allows us to study spheroid progression in a system where boundary conditions are negligible. We observe increased proliferation in E-cad+ spheroids and an inverse relationship between proliferation rate and collagen concentration. Our results demonstrate limited cell invasion at high collagen concentrations, which is consistent with the expected behavior based on the mechanical properties of the collagen gel [19, 21]. We then integrate system parameters and *in vitro* data into a computational continuum model. With this model, we analyze the influence of tumor system variables such as spheroid stiffness, cell polarization, proliferation rate and cell-ECM friction on tumor progression from a continuum mechanics perspective. Combining wet and dry experiments, we determine that matrix pore size and cell-matrix adhesion, both of which influence cell-ECM friction, are the mechanical features of a tumor with the greatest influence on tumor progression or suppression.

## 2. Results

### 2.1. Cancer cell proliferation is promoted by E-cad and limited by extracellular collagen

This work explores the influence of intercellular and extracellular mechanical conditions on the proliferation and invasion dynamics of tumor spheroids. These two main variables are chosen to modulate mechanical system conditions. Intercellular mechanics are controlled by the formation of cell-cell adherens junctions via expression of adhesion molecule E-cad. To this end, we study MDA-MB-231 breast cancer cells, which do not endogenously express E-cad (E-cad-), and MDA-MB-231 cells with E-cad lentiviral knock-in (E-cad+) [22]. We control the mechanical properties of the extracellular matrix (ECM) by seeding spheroids in collagen I gels at concentrations between 1 and 6 mg/mL. Our spheroid system utilizes the previously reported oil-in-water droplet microtechnology [23] to enclose E-cad-or E-cad+ MDA-MB-231 cells in an inner collagen I compartment suspended in a 100x larger collagen I matrix of the same concentration (Fig. 1a). With this system, we can simulate the collagen-rich stroma surrounding solid breast tumors [24] and analyze the changes in ECM mechanical properties as additional collagen is deposited during disease progression [18].

**Figure 1.**
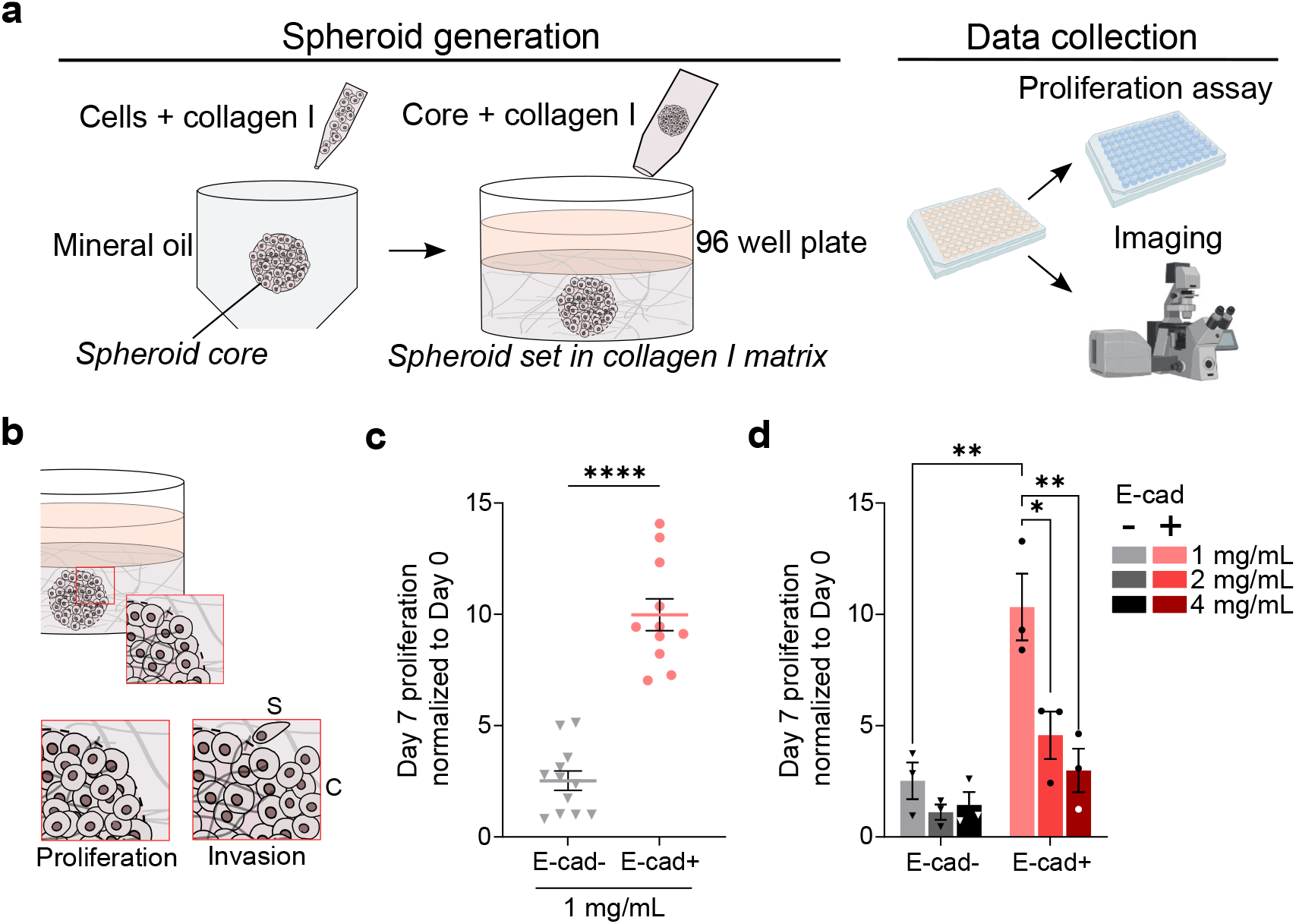
E-cad expression and ECM composition determine proliferation dynamics in spheroids. (a) Spheroid generation and data collection schematic. (b) Theoretical representation of the proliferation and invasion dynamics within tumor spheroids. S denotes single/detached cell invasion, C denotes collective invasion. (c) PrestoBlue relative proliferation on day 7 of MDA-MB-231 E-cad-/+ spheroids at 1 mg/mL extracellular collagen concentration. Data are mean ± SEM. Statistical test used: unpaired t-test, ****P ≤ 0.0001. (d) PrestoBlue relative proliferation of MDA-MB-231 E-cad-/+ spheroids at extracellular collagen concentration of 1, 2 and 4 mg/mL comparing spheroids on day of generation (day 0) to day 7. Data are mean ± SEM. Statistical test used: two-way ANOVA, **P ≤ 0.01, *P ≤ 0.05. The data in (c) and (d) are reported for N=3 biological replicates and n=3+ technical replicates.

E-cad expression modulates intercellular mechanics through adherens junctions that stiffen the cell aggregate to favor collective cell invasion and promotes increased proliferation dynamics of the cellular system (Fig. 1b). E-cad is a transmembrane protein that is ubiquitously expressed in non-invasive epithelial and tumor cells prior to their epithelial-mesenchymal transition (EMT) [19]. For this reason, it was believed to be a tumor-suppressor gene, and its loss a prerequisite for tumor metastasis; however, this hypothesis has been recently questioned [25, 26, 27]. We showed that E-cad expression results in hyper-proliferation of breast cancer cells by promoting activation of the ERK cascade utilizing a similar 3D organoid model. In our spheroids, we observe an increase in proliferation in E-cad+ spheroids compared to E-cad-of more than four-fold in the case of 1 mg/mL collagen matrices (Fig. 1c). This enhanced E-cad dependent proliferation is maintained at all collagen concentrations, supporting the previously reported hyper-proliferation observed in 3D cultures [27].

Regarding the effects of the ECM on cancer cell proliferation, we observe that softer matrices (1 mg/mL) promote higher proliferation rates than stiffer matrices regardless of E-cad expression (Fig. 1d). Specifically, we observed more than a three-fold decrease in E-cad+ spheroid proliferation at 4 mg/mL collagen concentration compared to the 1 mg/mL condition. Softer matrices sustained higher proliferation rates over time, while proliferation rates plateau over time in more rigid matrices (Fig. S1a). While many previous studies have reported increasing cell proliferation with the elastic modulus of the ECM, we observe decreasing proliferation with collagen concentration/elastic modulus [19]. These results may be explained by a reduction in the pore size of the ECM leading to higher frictional forces that, in turn, constrain spheroid spatial growth. As a consequence, cell proliferation is limited to a maximum local cell density that we define as the maximum carrying capacity of the spheroid. This maximum carrying capacity is determined by the consumption rate of nutrients, the spheroids’ spatial cell concentration, and the physical properties of the collagen matrix.

Cell invasion patterns also influence spheroid carrying capacity. As cells invade the collagen bulk, the local cell density is dispersed across the invaded area. Therefore, this proliferation dynamic cannot be analyzed independently, but requires a coupled analysis with cell invasion to understand the spatial constraints on spheroid growth and tumor progression. This phenomenon is difficult to prove experimentally but is later addressed with the help of a computational model.

Although both intercellular and extracellular variables have significant effects on proliferation, we observe that E-cad expression strongly correlates with proliferation as all E-cad+ spheroids surpass the relative proliferation observed in all E-cad-conditions on both day 5 and day 7 (Fig. 1d, Fig. S1a). It must be noted that changes in collagen concentration affect several ECM properties including elastic modulus, pore size, fiber alignment, and cell speed [19]. These parameters may have different and even contradicting effects on cancer cell proliferation. It is challenging to engineer matrices where each property can be tuned individually, thus it is difficult to experimentally uncouple their relative contributions to cancer cell proliferation. Therefore, we look to computational tools to evaluate the effects of each underlying mechanism individually.

### 2.2. Collective cell invasion is promoted by E-cad and limited by extracellular collagen

In addition to proliferation, another key feature of tumor progression is cell invasion. It not only dictates the fate of the tumor by means of spatial expansion and potential metastasis, but also intrinsically encourages proliferation. A clear consequence of collective invasion is the regulation of spatial confinement and access to nutrients, which determines cell survival within the spheroid. This relation between proliferation and invasion is correlative through the previously mentioned spheroid carrying capacity; an increase in invasion leads to mechanical stress within the spheroid, thus leading to radial forces projected from the spheroid that promote the expansion of the cell aggregate. Therefore, both responses are intrinsically coupled and highly dependent on both intercellular and extracellular conditions. In addition, when evaluating invasion, it is important to distinguish between collective and single-cell phenotypes (Fig. 1b). Intercellular adhesion promotes an orchestrated response of the system. Mechanical waves travel within the spheroid transmitting propulsion forces from the spheroid’s leading edge and compressive forces from the inner volume to the intermediate regions [28]. However, low intercellular adhesion hinders the transmission of such mechanical waves and encourages mesenchymal invasion from the outer (leading) surface of the spheroid. To better differentiate these two invasion phenotypes, we define within our spheroids a cellular collective region and a detached (mesenchymal) cell region (Fig. 2a).

**Figure 2.**
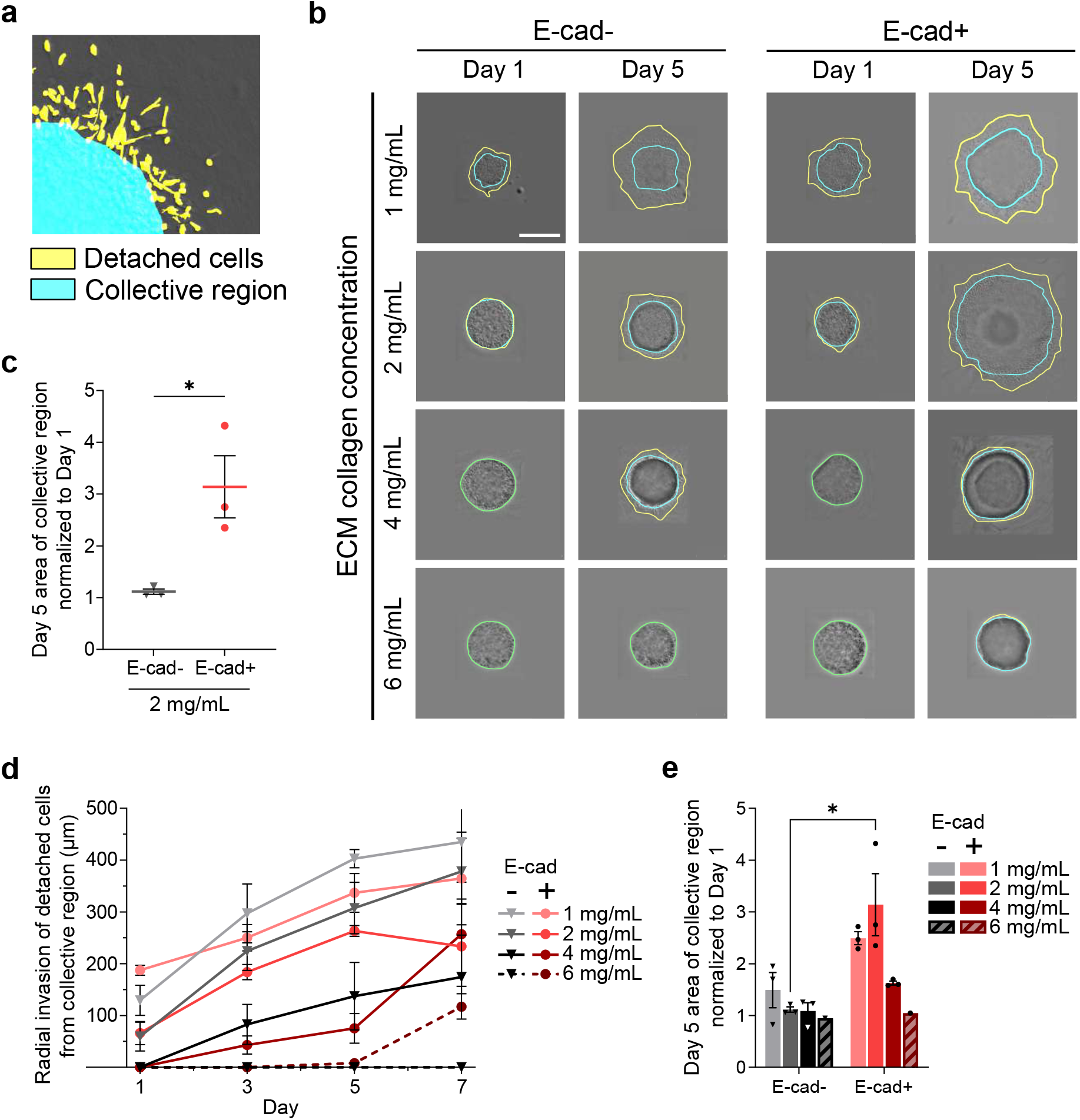
E-cad expression and ECM collagen concentration impact spheroid invasion. (a) Schematic defining the detached cells region in yellow and the collective region in cyan. (b) DIC microscopy images of MDA-MB-231 breast cancer cells (E-cad-lentiviral control, left panel) and MDA-MB-231 cells with E-cad lentiviral knock-in (E-cad+, right panel) on day 1 and day 5. Yellow annotations label the detached cells region and cyan annotations label the collective region. Scale bar: 1 mm. (c) E-cad dependent expansion of collective region area of tumor spheroids from day 1 to day 5 in a 2 mg/mL collagen ECM. Data are mean ± SEM. Statistical test used: unpaired t-test, *P ≤ 0.05. N = 3. (d) Average radial invasion distance from the edge of the collective region to the edge of the detached region over time for all E-cad conditions (-/+) and extracellular collagen concentrations (1, 2, 4, 6 mg/mL). The error bar for 1 mg/mL E-cad-day 7 is cut off to improve visuaization of all data. (e) Expansion of area of the collective region from day 1 to day 5 in E-cad-and E-cad+ spheroids at 1, 2, 4 and 6 mg/mL ECM collagen concentration. Data are mean ± SEM. Statistical test used: two-way ANOVA, *P ≤ 0.05. The data in (d) and (e) are reported for N=3 for 1, 2 and 4 mg/mL collagen concentrations. The data for 6 mg/mL are included as a limiting case with N=1 and n=3.

By annotating the boundary of the collective and detached cell regions, we determine both the extent and phenotype of cell invasion of each condition (Fig. 2b, Fig. S2). The results show E-cad+ spheroids invade the collagen bulk to a greater extent than E-cad-spheroids. The coupling between proliferation and invasion explains this difference. In E-cad+ conditions, we have observed that cells are hyperproliferative (Fig. 1c-d, Fig. S1a). Due to such proliferation within the spheroid, peripheral cells are pushed radially outward. These radial forces thus contribute to spheroid expansion to a larger spatial volume and allow further proliferation until a new critical cell density threshold is reached. Furthermore, the propagation efficiency of these mechanical waves is improved by E-cad expression. Adherens junctions formed in E-cad+ spheroids result in higher intercellular adhesion forces, and an increase in cell aggregate stiffness. This increased stiffness leads to higher mechanical stress and faster transmission speed within the spheroid because wave speed is directly related to mechanical stiffness [28]. The combined effects of E-cad expression are especially relevant within the collective region (Fig. 2c) where we observe increased collective invasion into the surrounding collagen ECM in E-cad+ conditions. In addition to proliferation-driven forces within the spheroid, the cells at the spheroid’s leading surface also exert traction forces. These forces arise from cells at the spheroid’s leading edge that, via focal adhesions, reorganize and pull on the surrounding collagen fibers as they invade. E-cad modulates this force by forming cell-ECM focal adhesions and intercellular adherens junctions. In E-cad+ conditions, the intercellular adherens junctions facilitate the transmission of traction forces to the dense internal region of the spheroid; however, if intercellular adhesion is low, as in E-cad-conditions, the cell-ECM propulsion forces are high enough to propel the peripheral cells outward from the spheroid. High propulsion forces result in single-cell invasion, and as a consequence, the traction force is not propagated to the core of the spheroid. The extent of single-cell invasion is quantified for each spheroid by measuring the average radial distance between the edge of the collective region and the leading invasion front (Fig. 2d). Under the same extracellular conditions, detached cells invade a larger radial distance from the collective region in E-cad-spheroids, i.e., a larger single-cell invaded area. These results confirm the invasion phenotypes that have been previously observed in E-cad-and E-cad+ cell lines: collective invasion is more often observed in E-cad+ cells while E-cad-cells invade as single cells with a mesenchymal phenotype [27].

While traction forces from cells at the spheroid’s leading edge and radial expansion forces due to proliferation support invasion into the ECM, friction opposes this invasion. Friction forces are determined by the mechanical and structural properties of the ECM; therefore, they are modulated in our spheroid system by the collagen concentration used. An increase in collagen concentration results in a smaller pore size and a change in the elastic modulus of the collagen gel, which controls the ability of cells to reorganize the matrix [19]. Therefore, the net friction force depends on collagen concentration. We observe decreases in the collective cell area at high collagen concentrations where the elastic modulus is high and the average pore size is low (Fig. 2e, Fig. S1b) [19]. In these conditions, high cell-ECM friction limits both collective and detached cell invasion despite the previously described tensile stresses due to proliferation and cell-cell or cell-ECM adhesion. In extreme cases (6 mg/mL collagen), null tumor expansion is observed. Spheroid confinement due to collagen concentration also explains the decrease in proliferation observed as collagen concentration increases (Fig. 1d, Fig. S1a) and reinforces our understanding of a limiting cell density threshold impeding further growth, i.e. spheroid carrying capacity. However, note that the increase in collagen concentration from 1 mg/mL to 2 mg/mL encourages collective invasion.

The average pore size of a collagen gel decreases with collagen concentration from 1 mg/mL to 2 mg/mL; however, the elastic modulus of the gel also decreases from 1 mg/mL to 2 mg/mL [19]. Cells can more easily remodel a gel with a lower elastic modulus to allow for continuum expansion. The increase in collagen fibers could also provide more potential ligands for cells to adhere to and propel through the matrix. We next evaluate this hypothesis by quantifying peripheral cell polarization, which indicates stronger propulsion forces at the spheroid’s boundary [29, 30].

### 2.3. Peripheral cell polarization is limited by E-cad and promoted by extracellular collagen

The ability of cells at the leading edge of the spheroid to adhere to surrounding collagen fibers and exert propulsion forces is in part dependent on the ability of cells to polarize [29, 30]. Thus, cell invasion into the surrounding collagen matrix is dependent on cell polarization. From a mechanical perspective, when a cell that is circular in shape (epithelial) contracts while using ECM fibers as support points, it generates a system of forces with radial symmetry so that the resultant force at its center of mass is null. This prevents the effective propulsion force that would be required for the cell to invade the matrix. If the cell is elongated (mesenchymal), radial symmetry is lost, and an effective force is generated along the direction of polarization [29, 31]. This effective force does not only depend on cell polarity but also on other features such as the formation of focal adhesions, the stiffness and viscous relaxation of the ECM matrix, and the active stress within the cell [29, 32, 30]. Therefore, the variations introduced in this work at the intercellular and extracellular levels may alter cell polarization, explaining in part the invasion patterns presented in Fig. 2.

To examine the effects of E-cad expression and collagen concentration on the ability of cells to generate the propulsion force required for invasion, we quantified the polarity of cells at the spheroid’s edge (Fig. 3a-b, Fig. S3). This analysis demonstrates that E-cad-cells are more polarized than E-cad+ cells (Fig. 3b-c, Fig. S1c). This inverse relationship of E-cad expression and polarity is explained by the decrease in intercellular adhesion forces when E-cad is not expressed, which allows cells at the spheroid front to polarize without the opposition of high traction stresses from the cell aggregate. The lack of E-cad expression results in highly polarized cells that invade as single/detached cells, as expected due to E-cad’s role in EMT [14] and the removed boundary constraints that can arise from surrounding cells. Alternatively, expression of E-cad (i.e., E-cad+) promotes cell-cell adhesion resulting in higher mechanical boundary constraints on the cells. These cells invade collectively and have lower polarization scores.

**Figure 3.**
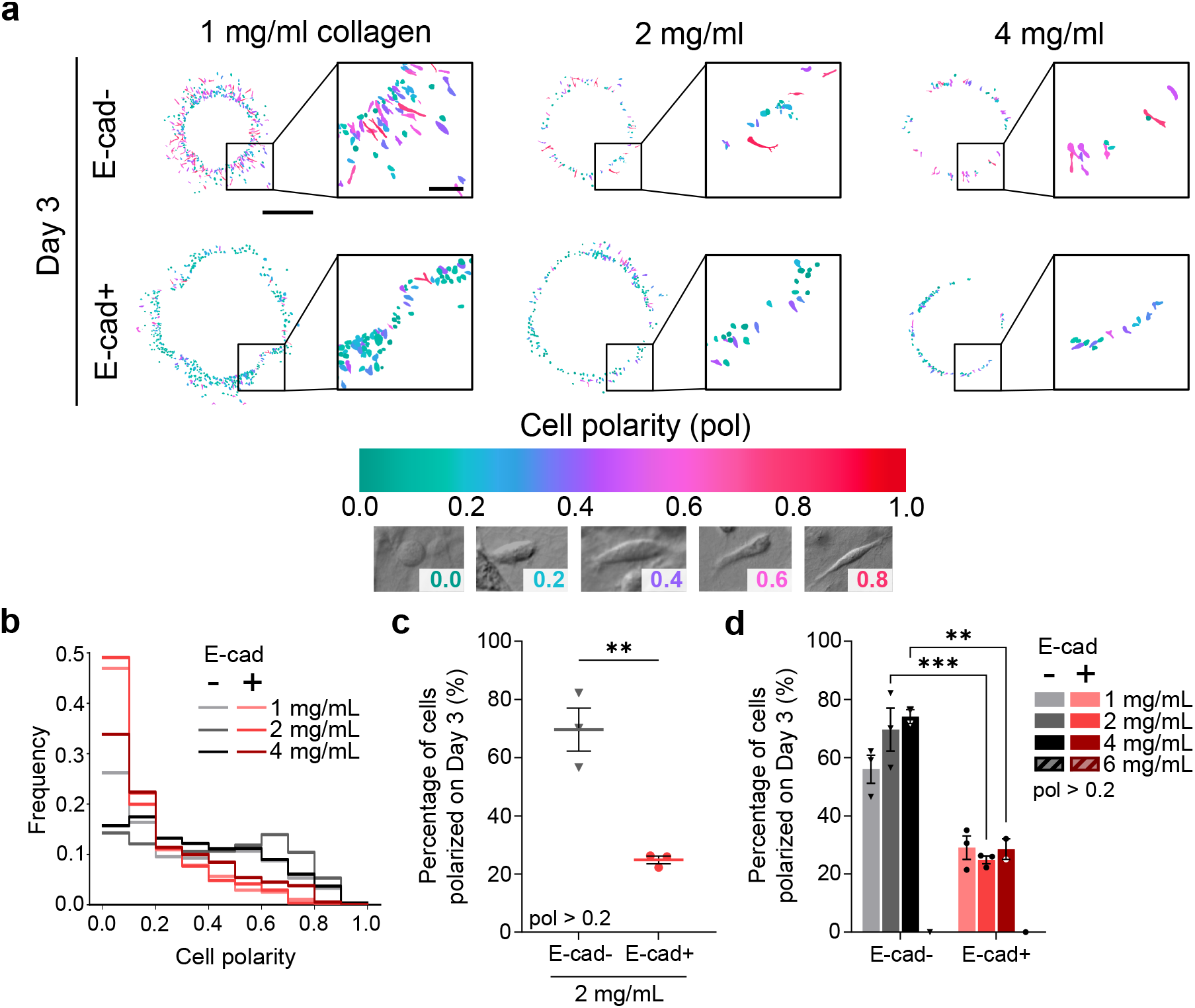
Cell polarity is influenced by E-cad expression and ECM concentration. (a) Morphology masks of cells detached from spheroids on day 3, where cell polarity is quantified and correlated to mask color. Examples of cells with polarity 0.0, 0.2, 0.4, 0.6, 0.8 and 0.9. Scale bar: 1 mm, inset: 250 *μm.* (b) Relative frequency of cell polarities in different E-cad and matrix conditions. Cumulative values calculated over days 1, 3, 5 and 7 are shown. (c) Percentage of polarized/elongated cells (defined as having a cell polarity greater than 0.2) within the detached region by E-cad expression. Data shown are from day 3 in 2 mg/mL extracellular collagen. Data are mean ± SEM. Statistical test used: unpaired t-test, **P ≤ 0.01. N = 3. (d) Percentage of cells with polarity ¿ 0.2 on day 3 by extracellular collagen concentration. Data are mean ± SEM. Statistical test used: two-way ANOVA, *** P ≤ 0.001, **P ≤ 0.01. The data in (b) and (d) are from N = 3 (1 and 2 mg/mL) or N = 2 (4 mg/mL), n = 2-4. The data for 6 mg/mL collagen in (b) are included as a limiting case with N = 1 and n = 3. Each technical replicate analyzed ¿100 cells.

In addition to intercellular dependencies, the extracellular conditions (i.e., collagen concentration) are found to also influence cell polarization. Cell polarity increases with increasing collagen concentration (Fig. 3d, Fig. S1d). It must be noted that cell polarization results from a mechanical balance between traction forces from the spheroid and the surrounding ECM. Therefore, the stiffness and pore size of the ECM will significantly impact this mechanical balance by directing the resulting forces and modulating their magnitude. Thus, a higher collagen concentration reduces ECM pore size and increases the spatial density of collagen fibers that serve as support points for forces exerted by the cells [19]. The reduction in pore size forces the cells to adopt more elongated shapes to invade the ECM. In addition, the associated increase in stiffness [19] and presence of fibers affect the magnitude of the propulsion forces promoting invasion.

These polarization dynamics can explain the low collective invasion observed in collagen concentrations of 1 mg/mL (Fig. 2e). Despite lower frictional forces and larger spheroid proliferation, resulting in larger pushing forces from the internal region of the spheroid, the propulsion forces are smaller as a consequence of low polarity (Fig. 3b). These reduced propulsion forces generated at the leading edge of the spheroid are not sufficient for the cells to collectively invade the collagen matrix. Detached cell invasion requires smaller propulsion forces, so these polarization effects are not as prevalent in the single-cell invasion results (Fig. 2d).

### 2.4. A mechanistically based theoretical model explains proliferation and invasion dynamics

The experimental results from these tumor spheroids demonstrate the importance of both intercellular and extracellular mechanical conditions on proliferation and invasion. An overexpression of E-cad is found to promote proliferation and hinder cell polarization at the leading edge of the spheroid. Moreover, an increase in extracellular collagen concentration is found to reduce proliferation and hinder collective invasion. However, this experimental analysis is limited to systematic correlations and does not allow for a deeper identification of the specific mechanobiological properties governing such processes.

To provide more specific insights into the underlying mechanistic features governing proliferation and invasion, we present a physical model formulated at the continuum scale. The main physical biomechanisms described in the model are schematically presented in Fig. 4a. Cancer cells are conceived as a continuum cell aggregate to assess the spatial distribution of both proliferation and collective cell invasion [33]. Here, we define the cell aggregate as a continuum where local mechanical stimuli at the cellular level can be globally transmitted in the form of mechanical waves and thus can transport interaction forces within the cell continuum [34, 35, 36, 37]. In parallel to these mechanical waves, local sources of cell proliferation lead to increases in cell density, resulting in internal stresses within the continuum. These internal stresses oppose the external pressure from the ECM. The mechanical balance between such internal and external forces governs the expansion of the cell aggregate. Capturing a complete formulation of cancer cell proliferation and invasion requires both the mechanical balance and the spatial evolution of cancer cell density (see Methods for details on the formulation). The variables of the problem are the displacement and cell density fields. The main parameters of the problem, along with their link to proliferation and invasion, are outlined in Table 1.

**Figure 4.**
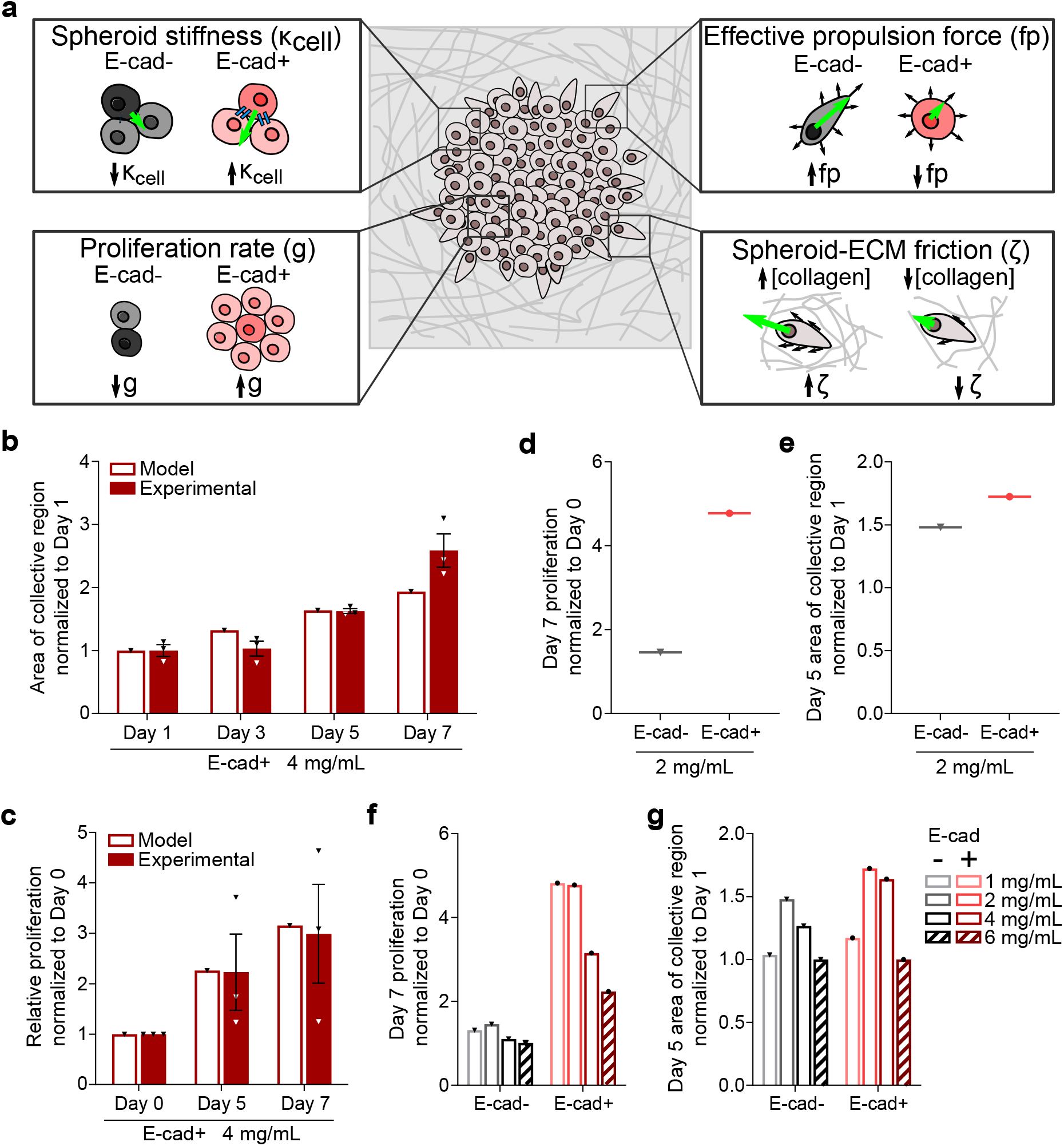
Modeling interpretation of spheroid mechanobiology from a continuum perspective. (a) Conceptual basis of the theoretical model representing a continuum spheroid embedded in a collagen matrix. The model accounts for four main biomechanical mechanisms, which are schematically described: the cell aggregate stiffness as a result of cell-cell adhesions (*κ_cell_*, top left); the proliferation rate associated to E-cad expression (***g***, bottom left); the effective propulsion force dependent on cell polarity (*f**p***, top right); and spheroid-ECM friction (ζ, bottom right). (b) Experimental invasion data and model predictions for the reference case (E-cad+ and 4 mg/mL collagen concentration). Experimental data presented as solid fill and model predictions presented as outline. (c) Experimental proliferation data and model predictions for the reference case. Experimental data presented in (b) and (c) are mean ± SEM. N = 3. E-cad dependence demonstrated in model results at 2 mg/mL extracellular collagen for (d) proliferation on day 7 and (e) continuum invasion on day 5. Model predictions of the influence of extracellular collagen concentration on (f) proliferation on day 7 and (g) continuum invasion on day 5.

**Table 1:** Key parameters in spheroid proliferation and invasion dynamics.

| Variable | Definition |
| --- | --- |
| Maximum carrying capacity ($\rho_\infty$) | The limiting cell density in a confined space preventing further proliferation due to overrate of nutrients consumption and lack of physical space. This parameter is difficult to evaluate experimentally and, therefore, an important outcome of the modeling framework. |
| Proliferation rate ($g$) | The proliferation rate of a single cell, which is influenced by E-cad expression. Note that the proliferation dynamics are not only determined by this variable but are also influenced by collective cell invasion and its modulation by the maximum carrying capacity. |
| Spheroid-ECM friction ($\zeta$) | The three-dimensional friction experienced by cells when invading through the ECM. Therefore, it strongly depends on the pore size of the collagen network that is determined by collagen concentration. |
| Spheroid stiffness ($\kappa_{cell}$) | The collective stiffness of the spheroid. Under compression, it represents the cell stiffness. Under tensile forces, it represents the mechanical resistance that a cell experiences when it tries to separate from the cell-aggregate. E-cad expression determines the formation of adherens junctions and intermediate actin filaments, therefore influences the structural integrity of the spheroid as a continuum and modulates this parameter. |
| Effective propulsion force ($f\mathbf{p}$) | The effective force exerted by the cells to invade. Such a cellular force is defined in a vector format with both a magnitude and direction. The magnitude is determined by the ability of the cells at the spheroid surface to propel themselves from the collagen fibers, dragging other cells from the interior of the spheroid. Therefore, it is influenced by the extracellular collagen concentration. Cell polarity determines how effective this force is. When the leading cells are highly polarized, cells invade more efficiently. The cell polarity can be obtained from the experimental results presented in Fig. 3 and depends on both E-cad expression and collagen concentration. |

The theoretical model serves as a virtual framework to combine all the mechanobiological parameters identified from the experiments and evaluate our hypotheses on each parameter’s influence on the proliferation and invasion dynamics of tumor spheroids in 3D collagen matrices. It is important to note that our experimental methodology controls E-cad expression and collagen concentration; however, these variables introduce a combination of biomechanical consequences with impacts on proliferation and invasion that cannot be isolated. The use of our modeling framework allows us to isolate each underlying mechanism and evaluate its independent role in modulating proliferation and invasion.

To this end, we first calibrate the model parameters by taking as reference an E-cad+ spheroid with a collagen concentration of 4 mg/mL. We choose this condition as a reference because large frictional terms limit invasion and allow us to determine the maximum carrying capacity (*ρ_∞_*). Further information on the motivation and values for the model parameters are detailed in Methods. Model predictions show a strong agreement with experimental results for both invasion (Fig. 4b) and proliferation (Fig. 4c). Then, we test the model dynamics by introducing relative variations to model parameters reflective of our experimental conditions (E-cad expression and collagen concentration) without further calibration (see Methods). Overall, the model results are in good agreement with the experimental results presented in Fig. 1 and Fig. 2 (see the comparison between experimental data and model predictions for all the conditions tested in Fig. S4a-b). Generally, an increase in proliferation rate associated with E-cad expression leads to higher cell densities and invasion (Fig. 4d-e). Alternatively, an increase in collagen concentration leads to lower invasion and a significant reduction in proliferation (Fig. 4f-g, Fig. S1e-f). A collagen concentration of 1 mg/mL is the condition where model predictions present higher deviations from experimental results.

### 2.5. Parametric analysis of the continuum model

Although the model predictions are in good agreement with experimental data for most conditions, the results for 1 mg/mL collagen concentration present higher deviations. Previous studies have reported that the mechanical properties (pore size, elastic modulus, etc.) of collagen gels are not linearly correlated with collagen concentration [19]. Thus, we anticipate a complicated relationship between collagen gel concentration and spheroid dynamics. We have confirmed this via inconsistent trends in our proliferation and invasion data as collagen concentration increases (Fig. 1d, Fig. 2e). To address these points, we provide a parametric study of the influence of each biomechanical parameter (spheroid/cell aggregate stiffness, proliferation rate, effective propulsion force, and spheroid-ECM friction, as depicted in Fig. 4a) on proliferation and invasion. Here, we first test how extracellular conditions impact spheroid growth. The effective propulsion force of the spheroid is the result of cell interactions with collagen fibers, thus we consider this force to be extracellular. In our parametric analysis, we present proliferation predictions as relative cell counts and invasion predictions as relative continuum radii. A reduction in propulsion force decreases both cell count and continuum radius (Fig. 5a). The opposite effect is observed for a reduction in cell-ECM friction, which increases cell count and continuum radius (Fig. 5b). When modulating propulsion force or spheroid-ECM friction, altering extracellular conditions has a greater effect on collective invasion than spheroid proliferation.

**Figure 5.**
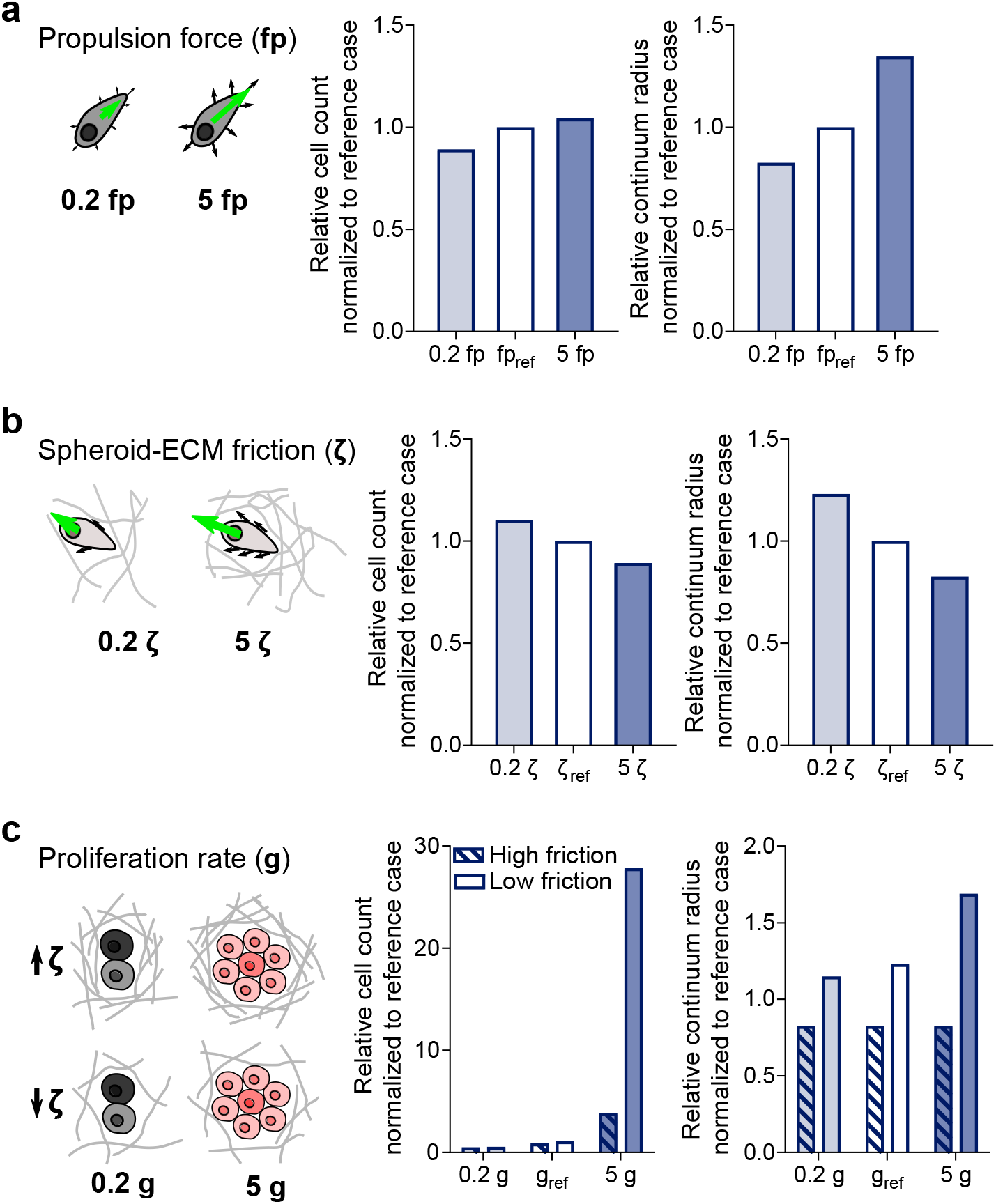
Parametric analysis of the continuum model with perturbations to extracellular and intercellular biomechanical properties. Model predictions of spheroid cell count and continuum radius when modulating the magnitude of extracellular conditions: (a) propulsion force or (b) spheroid-ECM friction. (c) Model predictions of spheroid cell count and continuum radius when modulating the magnitude of proliferation rate under high (5\**ζ_ref_*) and low (0.2\**ζ_ref_*) spheroid-ECM friction. All results shown are normalized to the reference case used in the simulations: E-cad+ in 4 mg/mL collagen matrix. All mechanical parameters are modified by dividing and multiplying the reference values by 5.

We consider proliferation rate an intercellular condition in our spheroid system due to its dependence on E-cad expression and the requirement for formation of cell-cell adherens junctions to activate the ERK hyperproliferation cascade [27]. Therefore, to analyze the impact of intercellular conditions, we alter the stiffness of the cell aggregate and the proliferation rate, which are both modulated experimentally by E-cad expression. When adjusting the spheroid stiffness, we predict minimal changes in cell count and continuum radius (Fig. S5). We then consider the proliferation rate under high and low cell-ECM friction (Fig. 5c). When altering the proliferation rate under conditions of high friction, we predict a moderate change in cell count and null variation in collective invasion. This suggests an important regulatory effect of cell-matrix friction on proliferation. In conditions where cell-matrix friction inhibits spatial expansion of the spheroid, the maximum carrying capacity (i.e., allowable cell density) of the system is quickly reached and cell proliferation is restricted. Under conditions of low cell-ECM friction, an increase in proliferation rate results in a significant change in both cell count and spheroid volume. This parametric analysis agrees with our experimental observations while identifying the influence of individual biomechanical parameters on spheroid proliferation and continuum invasion that cannot be isolated experimentally.

## 3. Discussion

Proliferation and invasion are two hallmarks of tumor progression that are often studied as independent processes despite their inherently cooperative nature. The molecular mechanisms and biomechanical features of these two processes are generally investigated as mutually exclusive events, according to the “go or grow” hypothesis [3, 4, 5]. However, from a clinical perspective, this overlap is recognized in both breast cancer diagnosis and standard of care treatments. One of the key determinants in breast cancer staging is tumor size, which is the result of combined tumor cell proliferation and continuum invasion at the primary site [38]. Non-surgical or neoadjuvant therapies often include chemotherapy or radiation, both of which aim to shrink the primary tumor size and kill cancer cells systemically [39, 40]. While clinical practice for breast cancer appears to account for the intrinsic coupling of proliferation and invasion of a primary tumor, scientific research often continues to study these processes independently [6, 7, 8, 9].

Here, we first demonstrate that proliferation and invasion are intrinsically coupled using 3D experimental assays on breast cancer spheroids modulating two main variables: expression of intercellular adhesion protein E-cad and extracellular collagen content. These data show that E-cad expression encourages both proliferation and collective invasion, while increases in ECM collagen concentration generally hinder both processes. From a mechanobiology perspective, E-cad expression promotes cell-cell adhesion that hinders cell polarization and, a priori, should hinder cell invasion. However, E-cad expression increases cell proliferation [27]. This hyper-proliferation leads to an increase in tumor spheroid volume and, therefore, a larger continuum invaded area. Increases in collagen (secreted by tumor and stromal cells) change the pore size and elastic modulus of the matrix [18, 19], leading to higher frictional forces opposing invasion. This reduction in invasion is accompanied by a reduction in proliferation as well.

Manipulation of E-cad expression and collagen concentration leads to the modulation of multiple biomechanical parameters; however, each mechanism’s influence on proliferation and invasion cannot be isolated experimentally. Limited collective invasion may be in part due to a low proliferation rate that is not sufficient to support spatial expansion of the spheroid, and vice versa. Thus, we establish a continuum mechanical model to delineate the impacts of individual biomechanics on proliferation and invasion dynamics.

Our continuum model provides insights into how E-cad expression and collagen concentration change specific biomechanical parameters and the consequences of those changes. Both extracellular (propulsion force, cell-ECM friction) and intercellular (E-cad mediated proliferation rate) mechanics are further explored independently through this model, explaining experimental observations. For example, our experimental data demonstrates that E-cad expression increases proliferation rate and cell-cell adhesion in the spheroid continuum, but decreases cell polarity. The increase in proliferation rate induces compression stresses within the spheroid encouraging continuum expansion. The increase in cell-cell adhesion induces a higher mechanical resistance to single-cell invasion (i.e., spheroid deformation from a continuum perspective). The decrease in cell polarity reduces the effective propulsion force and, therefore, continuum invasion. However, extracellular factors such as collagen concentration impact cell-ECM friction and cell polarity as well. When collagen concentration increases, friction and polarity also increase. The resulting system balance is a competition between propulsion forces, which increase with cell polarity, and increased friction. Both model and experimental results demonstrate a predominant role of friction forces in determining the proliferative and invasive fate of the tumor spheroid. In our system, these friction forces are modulated by E-cad-dependent cell-ECM adhesion and matrix pore size.

In one experimental condition (the lowest probed collagen concentration, 1 mg/mL), experimental results differ from model predictions. The divergence is likely due to the nonlinear relationship of certain collagen gel features that impact cell-ECM friction, further demonstrating how impactful friction is on tumor fate. For example, the average pore size for a 1 mg/mL collagen gel is significantly larger than the pore size of 2 mg/mL and 4 mg/mL gels [19]; therefore, cell-ECM friction is reduced in the 1 mg/mL condition. Low friction means there is a reduced resistance to collective invasion, and the spheroid can expand, thus accommodating cell proliferation and increasing the system’s carrying capacity. In this special case, friction forces are overcome by the compression stresses and expansion forces promoting spheroid growth. However, low collagen concentration and large pore size also decrease the frequency and strength of cell-ECM adhesion thus reducing the effective propulsion forces and resulting invasion of the continuum into the matrix. We conclude that a reduction in propulsion forces and cell-ECM friction due to the low collagen concentration in the 1 mg/mL condition encourages proliferation but limits continuum invasion (Fig. 5). This explanation reconciles the disagreement between experimentally measured invasion and model predictions of invasion while also supporting the agreement in proliferation results.

It is important to emphasize that the theoretical framework presented is conceptualized from a continuum basis and aims to model the proliferation and invasion dynamics from a cell aggregate (collective) rather than a single-cell (detached) perspective. We chose this approach to best study the proliferation and invasion dynamics in E-cad+ cancers, which report decreased survival compared to E-cad-cancers specifically in invasive ductal carcinomas [17]. In the region of the matrix where cells are detached, the cellular system cannot be treated as a continuum, and alternative models should be explored, such as single-cell invasion models [41, 42, 43, 44], mechanistic protrusive-based models [45, 46, 47], motor-clutch based models [32, 48] or multiscale approaches [49, 50]. Regarding the continuum description of the problem, further efforts should focus on incorporating propulsion force dependencies with pore size, which we have identified as the main contributor to spheroid dynamics at all extracellular concentrations examined.

Overall, we demonstrate the advantages of complementary experimental and computational methodologies to address mechanobiological elements of tumor progression that are difficult to isolate experimentally. We provide a virtual testbed for the analysis of isolated biomechanical parameters governing proliferation and invasion of breast cancer spheroids, and we use this platform to identify cell-ECM friction as a key regulator of tumor progression driven by intercellular (E-cad dependent cell-ECM adhesion) and extracellular (matrix pore size) properties of the tumor microenvironment.

## 4. Methods

### 4.1. Cell culture

MDA-MB-231 shRNA transfection control (E-cad -) and E-cadherin lentiviral knock-in (E-cad +) previously described by Lee et al. [22] were maintained in Dulbecco’s modified Eagle’s medium (DMEM, Corning, 10-013-CV) supplemented with 10% (v/v) fetal bovine serum (FBS, Corning, 35-010-CV), 1% (v/v) Penicillin-Streptomycin (Gibco, 15140-122), and 5 *μg/ml* puromycin (Gibco, A11138-03). Cells were maintained at 37^°^C and 5% *CO_2_*

### 4.2. Spheroid culture

Spheroid cores were generated by the oil-in-water droplet technique previously described by Lee et al. [22] and 3D collagen I matrices following a modified version of the protocol described by Fraley et al. [19]. This model’s framework was adapted from Jimenez, et al. [51]. Briefly, collagen I was prepared by thoroughly mixing equal parts cell culture medium (Dulbecco’s modified Eagle’s medium supplemented with 10% fetal bovine serum and 1% Penicillin-Streptomycin) and reconstitution buffer (1.1 g sodium bicarbonate and 2.4 g HEPES in 50 ml milli-Q water), adding collagen I (Corning, HC, Rat Tail) to 1, 2, 4, or 6 mg/ml final concentration, and 1M NaOH at a 4% v:v ratio to the collagen volume. Cells were resuspended in collagen I at a density of 1E+04 cells/*μl* and 1 *μl* droplets were allowed to gel in oil columns for 1 hour at 37 ^o^C. Cores were resuspended in a bulk collagen I gel of the same concentration, and 100 *μl* of the mixture was plated in a preheated 96 wells glass bottom plate (Cellvis) with one core centrally located in each well. The spheroid was incubated at 37 ^o^C for 1 hour to allow the bulk collagen to gel before a warmed culture medium was added to match the total gel volume.

### 4.3. Spheroid imaging

Differential interference contrast (DIC) microscopy images of live spheroids were taken on days 1, 3, 5 and 7 using a Nikon A1R Confocal mounted on a Nikon Eclipse Ti inverted microscope and OKO Labs stage top incubator to control temperature and *CO_2_.* Images were taken with a 10X/0.30 Plan Fluor objective, N1 DIC condenser and a 10X DIC slider (Nikon). GFP (488 nm) or RFP (561.5 nm) lasers were run concurrently with transmitted light. Images were taken at 2048 resolution with 2X line averaging.

### 4.4. PrestoBlue viability assay

Spheroids were incubated in a 1X PrestoBlue (Invitrogen) solution diluted in normal culture medium for 3 hours at 37 ^o^C. Blank wells prepared with 1X PresotBlue were measured and subtracted from spheroid signals to account for background signal. A SpectraMax plate reader (Molecular Devices) was used to read the fluorescence intensity (Excitation 540, Emission 600, Cutoff 590). PrestoBlue readings on days 5 and 7 are reported as a fold increase from day of generation (day 0). Lee et al. has previously demonstrated that PrestoBlue readings linearly correlate to a cell count [22].

### 4.5. Image analysis

DIC microscopy images of the spheroids were corrected for illumination inhomogeneities and stitched using the ImageJ stitching plugin [52]. The inner collective and detached cell regions were manually segmented for all replicas. Migration distance from the collective region was defined as the mean of the minimum distance from all points in the outer edge of the detached region to the edge of the collective region.

Regarding cell polarity analysis, at least 100 cells were manually segmented, segmenting all cells in a spheroid region clockwise until at least 100 cells and a quarter of the spheroid had been segmented for each technical replicate. Cell circularity was defined as 4*π*(*Area/Perimeter*^2^) [53]. Cell polarity was defined as 1 — *cellcircularity*. A polarity value near zero means the cell presents a rounded morphology, whereas a value near one indicates that the cell is highly polarized. To calculate the percentage of polarized cells in each condition, cells were considered to be polarized when their polarity was above 0.2 [54, 55]. Normalized histograms of detached cells’ polarity were calculated for all conditions (expression or absence of E-cadherin, different extracellular collagen concentrations) on days 1, 3, 5, and 7 of spheroid growth. Cumulative histograms were calculated by accounting for all cells from the same conditions at all time points of the experiment. Image data analysis was performed using custom Python codes.

### 4.6. Continuum model

For the mechanical balance, we assume negligible inertial effects and, in its spatial form, reads as:

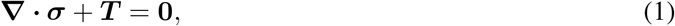

where **∇** is the spatial gradient. The mechanical stress within the cell aggregate is described, in its deformed state, by the Cauchy stress tensor ***σ***. The term ***T*** describes the external body forces.

The stress within the cell aggregate ***σ*** is defined by a constitutive equation which will depend on the specific cancer cells modeled. The external body mechanical sources to the cell aggregate (***T***) are mainly due to interactions between cells and the ECM [31, 29]. Therefore, it is convenient to split this contribution into a first component due to cell-ECM friction, and a second component related to propulsion forces exerted from the cell-ECM adhesion:

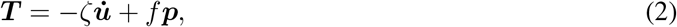

where *ζ* is the friction coefficient, 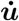 is the velocity field defined as the time derivative of the displacement field ***u***, ***p*** is the cell polarization, and *f* is the propulsion body force magnitude (per unit volume).

Moreover, we describe the evolution of local cell density *ρ* (here defined as number of cells per unit current volume) as the contribution of local proliferation sources, which depends on the maximum cell carrying capacity *ρ_∞_*, and a diffusion-like process mediated by an effective diffusion coefficient *d* as:

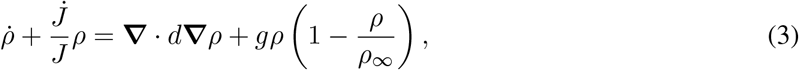

where *J* = *det* (***F***), with ***F*** being the deformation gradient. Note that the second term in the above expression is needed to account for the evolving spatial domain during proliferation, so that the mass balance is satisfied consistently [56, 57].

Note that the complete definition of the constitutive framework requires a constitutive equation for the cell aggregate stress (potentially linked to cell density) and the definition of the local proliferation source. Note also that the term *g* refers to the proliferation rate of cells per unit current volume.

The definition of the mechanical behavior of the cell aggregate is formulated to provide a direct link between the mechanical stress within the cells and the cell density by making use of a multiplicative decomposition of the deformation gradient ***F*** as [58, 59, 60]:

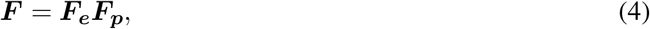

where ***F**_e_* and ***F**_p_* are the elastic and proliferation deformation gradient components, respectively. Note that the proliferation deformation is assumed volumetric and, therefore, the isochoric deformation (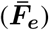) is only related to the elastic component as:

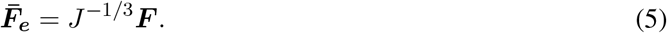

The energy potential describing the mechanical response of the cells is defined adopting a modeling scheme motivated by the neo-Hookean formulation as:

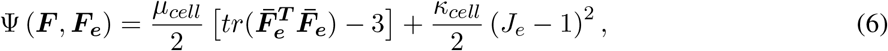

where *J_e_* = det (***F**_e_*) is the elastic Jacobian. The parameters *μ_cell_* and *κ_cell_* are, respectively, the apparent shear and bulk moduli of the cell aggregate describing the stiffness of the cell continuum to deform. Note that *κ_cell_* adopts different values depending on the stretch state. If *J_e_* < 1, the cells are locally under compression and this parameter refers to their material stiffness. Moreover, if *J_e_* > 1, this parameter is determined by the intercellular adhesion modulated by adhesive complexes and intermediate filaments. From thermodynamics principles the first Piola-Kirchhoff stress tensor can be derived as:

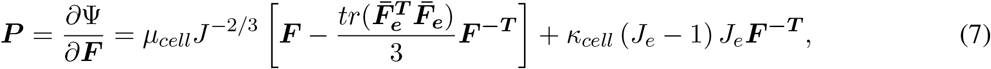

which is related to the Cauchy stress tensor by ***σ*** = *J*^-1^***PF**^T^*. Note that the proposed framework allows for the definition of other complex constitutive equations by simply changing the conceptualization of the energy potential Ψ. Thus, an additive composition of Ψ can be used to add viscoelastic components (see [61, 62] for consistent continuum visco-hyperelastic formulation).

Regarding the proliferation deformation gradient, it is directly linked to the current density as [63]:

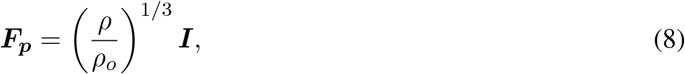

where ***I*** is the second order unit tensor and *ρ_o_* is the initial cell density.

### 4.7. Calibration of model parameters and motivations for perturbations

The model is conceptualized to isolate intercellular and extracellular mechanical properties in a simplified manner considering the cell aggregate as a continuum. We first calibrate the model parameters taking as reference a spheroid condition with overexpressed E-cad, i.e., E-cad+, and a collagen concentration of 4 mg/mL. In this condition, invasion is limited by large frictional forces, so the maximum carrying capacity (*ρ_∞_*) can be determined.

Then, all the variations in model parameters between experimental conditions are defined based on experimental observations without further fitting. In Table 2, we present the motivation for each choice.

**Table 2:** Motivation for parameter values for model calibration.

| Variable | Definition |
| --- | --- |
| Maximum carrying capacity ($\rho_\infty$) | This value is calibrated once with the reference case (E-cad+, 4 mg/ml) and kept constant for all conditions. |
| Proliferation rate ($g$) | This parameter differs from E-cad+ and E-cad- conditions and can be directly identified from the experimental data. Note that this parameter is kept constant for all extracellular conditions. |
| Spheroid-extracellular matrix friction ($\zeta$) | As discussed previously, this parameter is directly linked to the pore size of the collagen matrix. We first calibrate it for the reference case (4 mg/ml), as this parameter cannot be measured experimentally. Then, the friction coefficient for the different extracellular conditions is determined by an inverse relation with the pore volume of the surrounding collagen, so no fitting is used. This relation is taken from the experimental results obtained in a previous work by the authors [19]. This work provides the change in pore size depending on the collagen concentrations used herein. Therefore, the friction is found to be the highest at high collagen concentrations (6 mg/ml) and the lowest at low collagen concentrations (1 mg/ml) where the pores are larger. |
| Spheroid stiffness ($\kappa_{cell}$) | This parameter is related to the mechanical resistance of the spheroid. Under compressive forces, this parameter is defined as the cell stiffness [64, 65, 66]. In tensile conditions, it is defined as the force needed to detach cells. As adhesions between cells are the main cause of resistance to cell detachment, this parameter is defined by the E-cad expression of the cells. This stiffness is determined as the ratio of the mechanical stress prior to cell detachment to the corresponding strain. Note that this strain must be understood considering the two adhered cells as a continuum. The experimental data to obtain the values for E-cad- and E-cad+ conditions are taken from a previous work by the authors [15]. The stress is calculated as the force needed to separate two cells divided by the cross-sectional area of the cell. Such a force can be defined as the force of one E-cad/E-cad bond multiplied by the average number of bonds between two cells. Note that absent E-cadherin will reduce both the number of bonds and their strength. Therefore, the value for the spheroid stiffness in both E-cad+ and E-cad- conditions is directly taken from experimental data without *ad-hoc* fitting. |
| Effective propulsion force ($f_p$) | This parameter is defined by two variables: the propulsion body force and the cell polarity. The propulsion body force is kept constant for all conditions. This value is estimated considering that the force per individual cell is in the order of $10^2 - 10^3$ nN [67]. Therefore, the propulsion body force (i.e., per unit volume) in the spheroid can be computed as the force exerted by a single cell multiplied by the cell density (we have considered the initial cell density of our spheroids for this approximation). The cell polarity is directly taken from the experiments (Fig. 3, Fig. S1c-d), as the mean experimental value. Under E-cad+ conditions, where there is a strong coupling between cells, we assumed a linear distribution of the polarization to zero from the leading edge to the center of the spheroid. For E-cad- conditions, intercellular connections are weaker, which results in easier detachment of the leading cells and lower force transmission. For this case, the propulsion force was modeled as a steep exponential function where only cells very close to the edge have a non-negligible propulsion force. |

The model parameters used in the simulations are collected in Table S1. Note that the diffusion coefficient is kept constant to not influence the proliferation and migration dynamics between the tested conditions. This variable represents the tendency of cells to occupy regions of lower cell density.

## Acknowledgements

This work was supported through grants from the National Institute of Health (R01CA174388) to D.W., the National Cancer Institute (U54CA143868 and U54CA268083) to D.W., the National Institute of Arthritis and Musculoskeletal and Skin Diseases (U54AR081774) to D. W., and the National Institute on Aging (U01AG060903) to D.W. D.G.G, C.G.C and A.M.B. acknowledge support from the European Research Council (ERC) under the European Union’s Horizon 2020 Research and Innovation Program (Grant agreement No. 947723, project: 4D-BIOMAP). It was also supported by the Ministerio de Ciencia, Innovatión y Universidades, Agencia Estatal de Investigatión, under grant PID2019-109820RB-I00, MCIN/AEI/10.13039 /501100011033, cofinanced by European Regional Development Fund (ERDF), “A way of making Europe,” to A.M.B. C.G.C acknowledges support from the Ministerio de Universidades, Spain (FPU20/01459) and A.M.B. from Universidad Carlos III de Madrid (“Ayudas para la recualificación del profesorado funcionario o contratado”).

## Contributions

D.G.G. developed the hypothesis with input from G.C.R., D.W., and A.J.C experimentally, and A.M.B. computationally. A.J.C., G.C.R., and D.W. designed experiments. A.J.C. performed most experiments. G.C.R., W.H., and I.B. assisted with the experiments. C.G.C. performed most data analysis with input from D.G.G and A.M.B. A.J.C. assisted with data analysis. D.G.G. developed and implemented the computational model. C.G.C. assisted with the implementation of the computational model. D.G.G. and A.J.C. wrote the manuscript with input from C.G.C., G.C.R., D.W., and A.M.B.

## Ethics Declarations

The authors declare that they have no competing interests.

## References

[1] D. Hanahan, R. A. Weinberg, The hallmarks of cancer, Cell 100 (1) (2000) 57–70. doi:10.1016/S0092-8674(00)81683-9. URL https://doi.org/10.1016/S0092-8674(00)81683-9

[2] Y. A. Fouad, C. Aanei, Revisiting the hallmarks of cancer, Am J Cancer Res 7 (5) (2017) 1016–1036.

[3] D. Q. Matus, L. L. Lohmer, L. C. Kelley, A. J. Schindler, A. Q. Kohrman, M. Barkoulas, W. Zhang, Q. Chi, D. R. Sherwood, Invasive cell fate requires g1 cell-cycle arrest and histone deacetylase-mediated changes in gene expression, Developmental Cell 35 (2) (2015) 162–174. doi:https://doi.org/10.1016/j.devcel.2015.10.002. URL https://www.sciencedirect.com/science/article/pii/S1534580715006292

[4] C. R. Pfeifer, Y. Xia, K. Zhu, D. Liu, J. Irianto, V. M. M. García, L. M. S. Millán, B. Niese, S. Harding, D. Deviri, R. A. Greenberg, D. E. Discher, Constricted migration increases dna damage and independently represses cell cycle, Molecular Biology of the Cell 29 (16) (2018) 1948–1962, pMID: 29742017. arXiv:https://doi.org/10.1091/mbc.E18-02-0079, doi:10.1091/mbc.E18-02-0079. URL https://doi.org/10.1091/mbc.E18-02-0079

[5] K. S. Hoek, O. M. Eichhoff, N. C. Schlegel, U. Dobbeling, N. Kobert, L. Schaerer, S. Hemmi, R. Dummer, In vivo Switching of Human Melanoma Cells between Proliferative and Invasive States, Cancer Research 68 (3) (2008) 650–656. arXiv:https://aacrjournals.org/cancerres/article-pdf/68/3/650/2599834/650.pdf, doi:10.1158/0008-5472.CAN-07-2491. URL https://doi.org/10.1158/0008-5472.CAN-07-2491

[6] A. Kaznatcheev, J. G. Scott, D. Basanta, Edge effects in game-theoretic dynamics of spatially structured tumours, Journal of The Royal Society Interface 12 (108) (2015) 20150154. arXiv: https://royalsocietypublishing.org/doi/pdf/10.1098/rsif.2015.0154, doi:10.1098/rsif.2015.0154. URL https://royalsocietypublishing.org/doi/abs/10.1098/rsif.2015.0154

[7] P. Gerlee, S. Nelander, The impact of phenotypic switching on glioblastoma growth and invasion, PLOS Computational Biology 8 (6) (2012) 1–12. doi:10.1371/journal.pcbi.1002556. URL https://doi.org/10.1371/journal.pcbi.1002556

[8] H. Hatzikirou, D. Basanta, M. Simon, K. Schaller, A. Deutsch, ‘go or grow’: the key to the emergence of invasion in tumour progression?, Math Med Biol 29 (1) (2010) 49–65.

[9] J. A. Gallaher, J. S. Brown, A. R. A. Anderson, The impact of proliferation-migration tradeoffs on phenotypic evolution in cancer, Scientific Reports 9 (1) (2019) 2425. doi:10.1038/s41598-019-39636-x. URL https://doi.org/10.1038/s41598-019-39636-x

[10] T. Garay, Eva Juhasz, E. Molnár, M. Eisenbauer, A. Czirók, B. Dekan, V. László, M. A. Hoda, B. Dome, J. Tímár, W. Klepetko, W. Berger, B. Hegedus, Cell migration or cytokinesis and proliferation? - revisiting the “go or grow” hypothesis in cancer cells in vitro, Experimental Cell Research 319 (20) (2013) 3094–3103. doi:https://doi.org/10.1016/j.yexcr.2013.08.018. URL https://www.sciencedirect.com/science/article/pii/S0014482713003558

[11] C.-F. Gao, Q. Xie, Y.-L. Su, J. Koeman, S. K. Khoo, M. Gustafson, B. S. Knudsen, R. Hay, N. Shinomiya, G. F. V. Woude, Proliferation and invasion: plasticity in tumor cells, Proceedings of the National Academy of Sciences 102 (30) (2005) 10528–10533.

[12] E. Tuncer, R. R. Calçada, D. Zingg, S. Varum, P. Cheng, S. N. Freiberger, C.-X. Deng, I. Kleiter, M. P. Levesque, R. Dummer, et al., Smad signaling promotes melanoma metastasis independently of phenotype switching, The Journal of clinical investigation 129 (7) (2019) 2702–2716.

[13] J. P. Thiery, H. Acloque, R. Y. Huang, M. A. Nieto, Epithelial-mesenchymal transitions in development and disease, cell 139 (5) (2009) 871–890.

[14] K. Polyak, R. A. Weinberg, Transitions between epithelial and mesenchymal states: acquisition of malignant and stem cell traits, Nature Reviews Cancer 9 (4) (2009) 265–273.

[15] S. Bajpai, J. Correia, Y. Feng, J. Figueiredo, S. X. Sun, G. D. Longmore, G. Suriano, D. Wirtz, &#x3b1;-catenin mediates initial e-cadherin-dependent cell–cell recognition and subsequent bond strengthening, Proceedings of the National Academy of Sciences 105 (47) (2008) 18331–18336. arXiv:https://www.pnas.org/doi/pdf/10.1073/pnas.0806783105, doi:10.1073/pnas.0806783105. URL https://www.pnas.org/doi/abs/10.1073/pnas.0806783105

[16] G. C. Russo, A. J. Crawford, D. Clark, J. Cui, R. Carney, M. N. Karl, B. Su, B. Starich, T.-S. Lih, P. Kamat, Q. Zhang, P.-H. Wu, M.-H. Lee, H. S. Leong, V. W. Rebecca, H. Zhang, D. Wirtz, E-cadherin interacts with egfr resulting in hyper-activation of erk in multiple models of breast cancer, bioRxivarXiv:https://www.biorxiv.org/content/early/2022/01/25/2020.11.04.368746.full.pdf, doi:10.1101/2020.11.04.368746. URL https://www.biorxiv.org/content/early/2022/01/25/2020.11.04.368746

[17] C. Curtis, S. P. Shah, S.-F. Chin, G. Turashvili, O. M. Rueda, M. J. Dunning, D. Speed, A. G. Lynch, S. Samarajiwa, Y. Yuan, S. Gräf, G. Ha, G. Haffari, A. Bashashati, R. Russell, S. McKinney, C. Caldas, S. Aparicio, C. Curtis†, J. D. Brenton, I. Ellis, D. Huntsman, S. Pinder, A. Purushotham, L. Murphy, H. Bardwell, Z. Ding, L. Jones, B. Liu, I. Papatheodorou, S. J. Sammut, G. Wishart, S. Chia, K. Gelmon, C. Speers, P. Watson, R. Blamey, A. Green, D. Macmillan, E. Rakha, C. Gillett, A. Grigoriadis, E. de Rinaldis, A. Tutt, M. Parisien, S. Troup, D. Chan, C. Fielding, A.-T. Maia, S. McGuire, M. Osborne, S. M. Sayalero, I. Spiteri, J. Hadfield, L. Bell, K. Chow, N. Gale, M. Kovalik, Y. Ng, L. Prentice, S. Tavare, F. Markowetz, Y. Yuan, A. Langerød, E. Provenzano, A.-L. Børresen-Dale, M. E. T. A. B. R. I. C. Group, Co-chairs, W. committee, S. committee, Tissue, c. d. s. sites:, U. of Cambridge/Cancer Research UK Cambridge Research Institute, B. C. C. Agency, U. of Nottingham, K. C. London, M. I. of Cell Biology, C. g. c. centres:, D. a. subgroup:, The genomic and transcriptomic architecture of 2,000 breast tumours reveals novel subgroups, Nature 486 (7403) (2012) 346–352. doi:10.1038/nature10983. URL https://doi.org/10.1038/nature10983

[18] I. Acerbi, L. Cassereau, I. Dean, Q. Shi, A. Au, C. Park, Y. Y. Chen, J. Liphardt, E. S. Hwang, V. M. Weaver, Human breast cancer invasion and aggression correlates with ECM stiffening and immune cell infiltration, Integr Biol (Camb) 7 (10) (2015) 1120–1134.

[19] S. I. Fraley, P.-h. Wu, L. He, Y. Feng, R. Krisnamurthy, G. D. Longmore, D. Wirtz, Three-dimensional matrix fiber alignment modulates cell migration and mt1-mmp utility by spatially and temporally directing protrusions, Scientific Reports 5 (1) (2015) 14580. doi:10.1038/srep14580. URL https://doi.org/10.1038/srep14580

[20] M.-H. Lee, G. C. Russo, Y. S. Rahmanto, W. Du, A. J. Crawford, P.-H. Wu, D. Gilkes, A. Kiemen, T. Miyamoto, Y. Yu, et al., Multi-compartment tumor organoids, Materials Today.

[21] H. L. Hiraki, D. L. Matera, W. Y. Wang, E. S. Prabhu, Z. Zhang, F. Midekssa, A. E. Argento, J. M. Buschhaus, B. A. Humphries, G. D. Luker, et al., Fiber density and matrix stiffness modulate distinct cell migration modes in a 3d stroma mimetic composite hydrogel, Acta Biomaterialia.

[22] M.-H. Lee, P.-H. Wu, J. R. Staunton, R. Ros, G. D. Longmore, D. Wirtz, Mismatch in mechanical and adhesive properties induces pulsating cancer cell migration in epithelial monolayer, Biophysical Journal 102 (12) (2012) 2731–2741. doi:https://doi.org/10.1016/j.bpj.2012.05.005. URL https://www.sciencedirect.com/science/article/pii/S0006349512005577

[23] M.-H. Lee, Y. S. Rahmanto, G. Russo, P.-H. Wu, D. Gilkes, A. Kiemen, T. Miyamoto, Y. Yu, M. Habibi, I.-M. Shih, T.-L. Wang, D. Wirtz, Multi-compartment tumor organoids, bioRxivarXiv:https://www.biorxiv.org/content/early/2020/11/04/2020.11.03.367334.full.pdf, doi:10.1101/2020.11.03.367334. URL https://www.biorxiv.org/content/early/2020/11/04/2020.11.03.367334

[24] F. Kai, A. P. Drain, V. M. Weaver, The extracellular matrix modulates the metastatic journey, Developmental Cell 49 (3) (2019) 332–346. doi:10.1016/j.devcel.2019.03.026. URL https://doi.org/10.1016/j.devcel.2019.03.026

[25] K. Chu, K. M. Boley, R. Moraes, S. H. Barsky, F. M. Robertson, The paradox of e-cadherin: role in response to hypoxia in the tumor microenvironment and regulation of energy metabolism, Oncotarget 4(3) (2013) 446.

[26] V. Padmanaban, I. Krol, Y. Suhail, B. M. Szczerba, N. Aceto, J. S. Bader, A. J. Ewald, E-cadherin is required for metastasis in multiple models of breast cancer, Nature 573 (7774) (2019) 439–444.

[27] G. C. Russo, A. J. Crawford, D. Clark, J. Cui, R. Carney, M. N. Karl, B. Su, B. Starich, T.-S. Lih, P. Kamat, et al., E-cadherin interacts with egfr resulting in hyper-activation of erk in multiple models of breast cancer, BioRxiv (2022) 2020–11.

[28] N. Hino, L. Rossetti, A. Marín-Llaurads, K. Aoki, X. Trepat, M. Matsuda, T. Hirashima, Erk-mediated mechanochemical waves direct collective cell polarization, Developmental cell 53 (6) (2020) 646–660.

[29] R. Alert, X. Trepat, Physical models of collective cell migration, Annual Review of Condensed Matter Physics 11 (1) (2020) 77–101. arXiv:https://doi.org/10.1146/annurev-conmatphys-031218-013516, doi:10.1146/annurev-conmatphys-031218-013516. URL https://doi.org/10.1146/annurev-conmatphys-031218-013516

[30] F. Merino-Casallo, M. J. Gomez-Benito, S. Hervas-Raluy, J. M. Garcia-Aznar, Unravelling cell migration: defining movement from the cell surface, Cell Adhesion & Migration 16 (1) (2022) 25–64, pMID: 35499121. arXiv:https://doi.org/10.1080/19336918.2022.2055520, doi:10.1080/19336918.2022.2055520. URL https://doi.org/10.1080/19336918.2022.2055520

[31] D. Garcia-Gonzalez, A. Muñoz-Barrutia, Computational insights into the influence of substrate stiffness on collective cell migration, Extreme Mechanics Letters 40 (2020) 100928. doi:https://doi.org/10.1016/j.eml.2020.100928. URL https://www.sciencedirect.com/science/article/pii/S2352431620301826

[32] K. Adebowale, Z. Gong, J. C. Hou, K. M. Wisdom, D. Garbett, H.-p. Lee, S. Nam, T. Meyer, D. J. Odde, V. B. Shenoy, O. Chaudhuri, Enhanced substrate stress relaxation promotes filopodia-mediated cell migration, Nature Materials 20 (9) (2021) 1290–1299. doi:10.1038/s41563-021-00981-w. URL https://doi.org/10.1038/s41563-021-00981-w

[33] B. de Melo Quintela, S. Hervas-Raluy, J. M. Garcia-Aznar, D. Walker, K. Y. Wertheim, M. Viceconti, A theoretical analysis of the scale separation in a model to predict solid tumour growth, Journal of Theoretical Biology 547 (2022) 111173. doi:https://doi.org/10.1016/j.jtbi.2022.111173. URL https://www.sciencedirect.com/science/article/pii/S0022519322001710

[34] I. González-Valverde, J. M. García-Aznar, Mechanical modeling of collective cell migration: An agent-based and continuum material approach, Computer Methods in Applied Mechanics and Engineering 337 (2018) 246–262. doi:https://doi.org/10.1016/j.cma.2018.03.036. URL https://www.sciencedirect.com/science/article/pii/S0045782518301609

[35] R. Alert, C. Blanch-Mercader, J. Casademunt, Active fingering instability in tissue spreading, Phys. Rev. Lett. 122 (2019) 088104. doi:10.1103/PhysRevLett.122.088104. URL https://link.aps.org/doi/10.1103/PhysRevLett.122.088104

[36] S. Banerjee, M. C. Marchetti, Continuum Models of Collective Cell Migration, Springer International Publishing, Cham, 2019, pp. 45–66. doi:10.1007/978-3-030-17593-1_4. URL https://doi.org/10.1007/978-3-030-17593-1_4

[37] T. Bhattacharjee, D. B. Amchin, R. Alert, J. A. Ott, S. S. Datta, Chemotactic smoothing of collective migration, eLife 11 (2022) e71226. doi:10.7554/eLife.71226. URL https://doi.org/10.7554/eLife.71226

[38] A. C. Society, Stages of breast cancer: Understand breast cancer staging. URL https://www.cancer.org/cancer/breast-cancer/understanding-a-breast-cancer-diagnosis/stages-of-breast-cancer.html

[39] M. Clinic, Breast cancer (2022). URL https://www.mayoclinic.org/diseases-conditions/breast-cancer/diagnosis-treatment/

[40] A. C. Society, Breast cancer treatment: Treatment options for breast cancer. URL https://www.cancer.org/cancer/breast-cancer/treatment.html

[41] E. J. Campbell, P. Bagchi, A computational study of amoeboid motility in 3d: the role of extracellular matrix geometry, cell deformability, and cell-matrix adhesion, Biomechanics and Modeling in Mechanobiology 20 (1) (2021) 167–191. doi:10.1007/s10237-020-01376-7. URL https://doi.org/10.1007/s10237-020-01376-7

[42] A. Ippolito, V. S. Deshpande, Contact guidance via heterogeneity of substrate elasticity, Acta Biomaterialia doi:https://doi.org/10.1016/j.actbio.2021.11.024. URL https://www.sciencedirect.com/science/article/pii/S1742706121007728

[43] A. Ippolito, A. DeSimone, V. S. Deshpande, Contact guidance as a consequence of coupled morphological evolution and motility of adherent cells, Biomechanics and Modeling in Mechanobiology 21 (4) (2022) 1043–1065. doi:10.1007/s10237-022-01570-9. URL https://doi.org/10.1007/s10237-022-01570-9

[44] M.-C. Kim, Y. R. Silberberg, R. Abeyaratne, R. D. Kamm, H. H. Asada, Computational modeling of three-dimensional ecm-rigidity sensing to guide directed cell migration, Proceedings of the National Academy of Sciences 115 (3) (2018) E390–E399. arXiv:https://www.pnas.org/doi/pdf/10.1073/pnas.1717230115, doi:10.1073/pnas.1717230115. URL https://www.pnas.org/doi/abs/10.1073/pnas.1717230115

[45] B. M. Yeoman, P. Katira, A stochastic algorithm for accurately predicting path persistence of cells migrating in 3d matrix environments, PLOS ONE 13(11) (2018) 1–27. doi:10.1371/journal.pone.0207216. URL https://doi.org/10.1371/journal.pone.0207216

[46] F. Merino-Casallo, M. J. Gomez-Benito, R. Martinez-Cantin, J. M. Garcia-Aznar, A mechanistic protrusive-based model for 3d cell migration, European Journal of Cell Biology 101 (3) (2022) 151255. doi:https://doi.org/10.1016/j.ejcb.2022.151255. URL https://www.sciencedirect.com/science/article/pii/S0171933522000589

[47] I. G. Goncalves, J. M. Garcia-Aznar, Extracellular matrix density regulates the formation of tumour spheroids through cell migration, PLOS Computational Biology 17 (2) (2021) 1–22. doi:10.1371/journal.pcbi.1008764. URL https://doi.org/10.1371/journal.pcbi.1008764

[48] S. J. Tan, A. C. Chang, S. M. Anderson, C. M. Miller, L. S. Prahl, D. J. Odde, A. R. Dunn, Regulation and dynamics of force transmission at individual cell-matrix adhesion bonds, Science Advances 6 (20) (2020) eaax0317. arXiv:https://www.science.org/doi/pdf/10.1126/sciadv.aax0317, doi:10.1126/sciadv.aax0317. URL https://www.science.org/doi/abs/10.1126/sciadv.aax0317

[49] A. Buttenschon, L. Edelstein-Keshet, Bridging from single to collective cell migration: A review of models and links to experiments, PLOS Computational Biology 16 (12) (2020) 1–34. doi:10.1371/journal.pcbi.1008411. URL https://doi.org/10.1371/journal.pcbi.1008411

[50] B. A. Camley, W.-J. Rappel, Physical models of collective cell motility: from cell to tissue, Journal of Physics D: Applied Physics 50 (11) (2017) 113002. doi:10.1088/1361-6463/aa56fe. URL https://doi.org/10.1088/1361-6463/aa56fe

[51] A. M. Jimenez Valencia, P.-H. Wu, O. N. Yogurtcu, P. Rao, J. DiGiacomo, I. Godet, L. He, M.-H. Lee, D. Gilkes, S. X. Sun, D. Wirtz, Collective cancer cell invasion induced by coordinated contractile stresses, Oncotarget 6 (41) (2015) 43438–43451.

[52] S. Preibisch, S. Saalfeld, P. Tomancak, Globally optimal stitching of tiled 3d microscopic image acquisitions, Bioinformatics 25 (11) (2009) 1463–1465.

[53] N. I. of Health, Circularity. URL https://imagej.nih.gov/ij/plugins/circularity.html

[54] Y. L. Huang, C.-k. Tung, A. Zheng, B. J. Kim, M. Wu, Interstitial flows promote amoeboid over mesenchymal motility of breast cancer cells revealed by a three dimensional microfluidic model, Integrative Biology 7 (11) (2015) 1402–1411.

[55] M. I. Setyawati, C. Sevencan, B. H. Bay, J. Xie, Y. Zhang, P. Demokritou, D. T. Leong, Nano-tio2 drives epithelial-mesenchymal transition in intestinal epithelial cancer cells, Small 14 (30) (2018) 1800922.

[56] E. Comellas, S. Budday, J.-P. Pelteret, G. A. Holzapfel, P. Steinmann, Modeling the porous and viscous responses of human brain tissue behavior, Computer Methods in Applied Mechanics and Engineering 369 (2020) 113128. doi:https://doi.org/10.1016/j.cma.2020.113128. URL https://www.sciencedirect.com/science/article/pii/S0045782520303133

[57] M. Zarzor, S. Kaessmair, P. Steinmann, I. Blümcke, S. Budday, A two-field computational model couples cellular brain development with cortical folding, Brain Multiphysics 2 (2021) 100025. doi:https://doi.org/10.1016/j.brain.2021.100025. URL https://www.sciencedirect.com/science/article/pii/S2666522021000058

[58] S. Budday, P. Steinmann, On the influence of inhomogeneous stiffness and growth on mechanical instabilities in the developing brain, International Journal of Solids and Structures 132-133 (2018) 31–41. doi:https://doi.org/10.1016/j.ijsolstr.2017.08.010. URL https://www.sciencedirect.com/science/article/pii/S0020768317303669

[59] D. Ambrosi, M. Ben Amar, C. J. Cyron, A. DeSimone, A. Goriely, J. D. Humphrey, E. Kuhl, Growth and remodelling of living tissues: perspectives, challenges and opportunities, Journal of The Royal Society Interface 16 (157) (2019) 20190233. arXiv: https://royalsocietypublishing.org/doi/pdf/10.1098/rsif.2019.0233, doi:10.1098/rsif.2019.0233. URL https://royalsocietypublishing.org/doi/abs/10.1098/rsif.2019.0233

[60] E. Comellas, J. E. Farkas, G. Kleinberg, K. Lloyd, T. Mueller, T. J. Duerr, J. J. Munñz, J. R. Monaghan, S. J. Shefelbine, Local mechanical stimuli correlate with tissue growth in axolotl salamander joint morphogenesis, Proceedings of the Royal Society B: Biological Sciences 289 (1975) (2022) 20220621. arXiv:https://royalsocietypublishing.org/doi/pdf/10.1098/rspb.2022.0621, doi:10.1098/rspb.2022.0621. URL https://royalsocietypublishing.org/doi/abs/10.1098/rspb.2022.0621

[61] D. Garcia-Gonzalez, C. M. Landis, Magneto-diffusion-viscohyperelasticity for magneto-active hydrogels: Rate dependences across time scales, Journal of the Mechanics and Physics of Solids 139 (2020) 103934. doi:https://doi.org/10.1016/j.jmps.2020.103934. URL https://www.sciencedirect.com/science/article/pii/S0022509620301708

[62] C. Durcan, M. Hossain, G. Chagnon, D. Perić, L. Bsiesy, G. Karam, E. Girard, Experimental investigations of the human oesophagus: anisotropic properties of the embalmed muscular layer under large deformation, Biomechanics and Modeling in Mechanobiology 21 (4) (2022) 1169–1186. doi:10.1007/s10237-022-01583-4. URL https://doi.org/10.1007/s10237-022-01583-4

[63] D. Faghihi, X. Feng, E. A. Lima, J. T. Oden, T. E. Yankeelov, A coupled mass transport and deformation theory of multi-constituent tumor growth, Journal of the Mechanics and Physics of Solids 139 (2020) 103936. doi:https://doi.org/10.1016/j.jmps.2020.103936. URL https://www.sciencedirect.com/science/article/pii/S0022509620301721

[64] M. Nikkhah, J. S. Strobl, E. M. Schmelz, M. Agah, Evaluation of the influence of growth medium composition on cell elasticity, Journal of Biomechanics 44 (4) (2011) 762–766. doi:https://doi.org/10.1016/j.jbiomech.2010.11.002. URL https://www.sciencedirect.com/science/article/pii/S0021929010006159

[65] K. Pogoda, J. Jaczewska, J. Wiltowska-Zuber, O. Klymenko, K. Zuber, M. Fornal, M. Lekka, Depth-sensing analysis of cytoskeleton organization based on afm data, European Biophysics Journal 41 (1) (2012) 79–87. doi:10.1007/s00249-011-0761-9. URL https://doi.org/10.1007/s00249-011-0761-9

[66] K. Hayashi, M. Iwata, Stiffness of cancer cells measured with an afm indentation method, Journal of the Mechanical Behavior of Biomedical Materials 49 (2015) 105–111. doi:https://doi.org/10.1016/j.jmbbm.2015.04.030. URL https://www.sciencedirect.com/science/article/pii/S1751616115001587

[67] M. S. Hall, F. Alisafaei, E. Ban, X. Feng, C.-Y. Hui, V. B. Shenoy, M. Wu, Fibrous nonlinear elasticity enables positive mechanical feedback between cells and ecms, Proceedings of the National Academy of Sciences 113 (49) (2016) 14043–14048. arXiv:https://www.pnas.org/doi/pdf/10.1073/pnas.1613058113, doi:10.1073/pnas.1613058113. URL https://www.pnas.org/doi/abs/10.1073/pnas.1613058113

