## Supplementary Information for "Tumor proliferation and invasion are coupled through cell-extracellular matrix friction"

*Supplementary results*

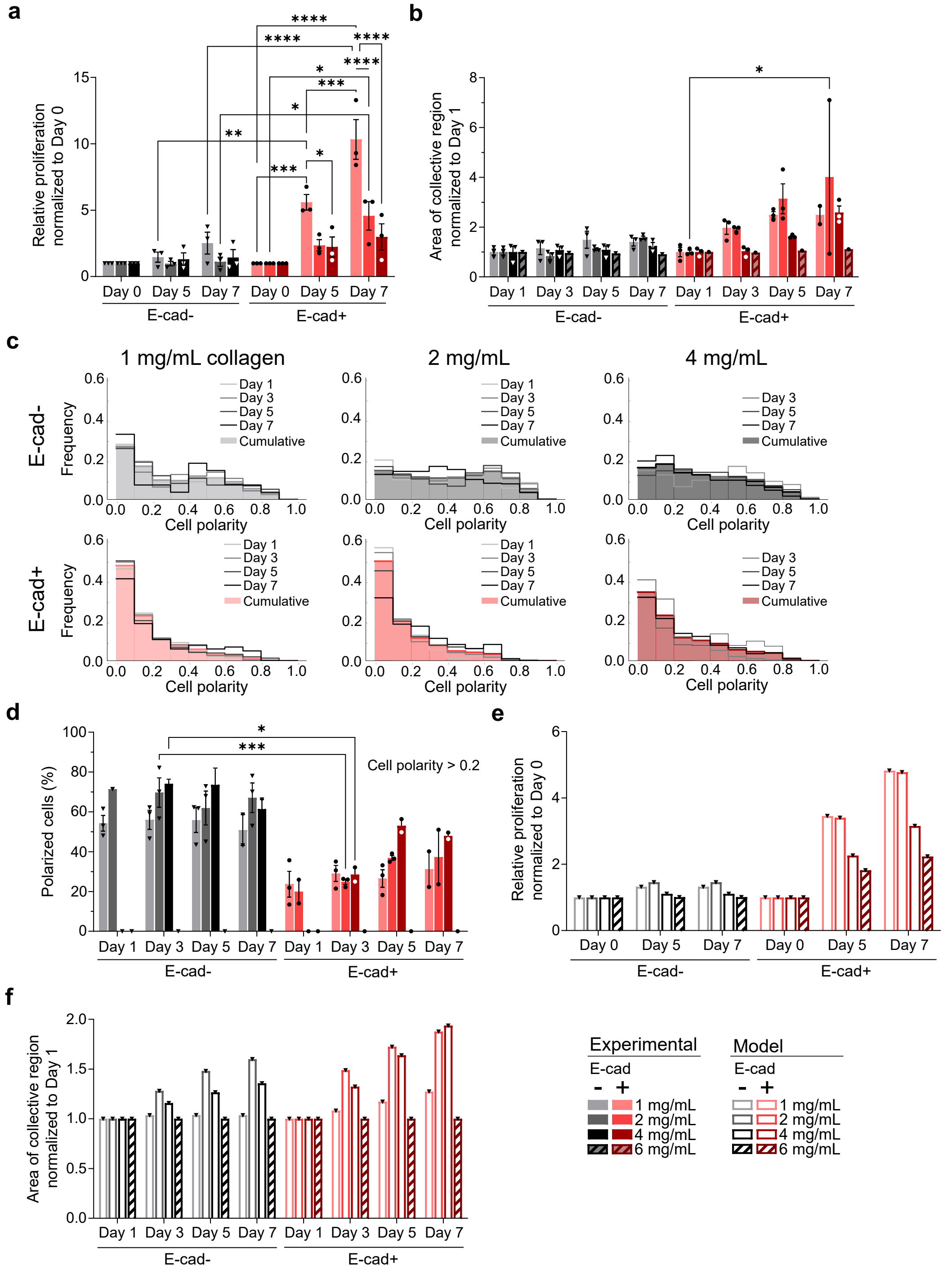


Figure S1. Proliferation, invasion, and polarization experimental data and model predictions by E-cad expression and collagen concentration at all time points. (a) PrestoBlue relative proliferation of MDA-MB-231 E-cad-/+ spheroids at extracellular collagen concentration of 1, 2 and 4 mg/mL comparing spheroids on day of generation (day 0) to days 5 and 7. (b) Expansion of area of the collective region from day 1 to days 3, 5 and 7 in E-cad-/+ spheroids at 1, 2, 4 and 6 mg/mL ECM collagen concentration collagen. (c) Relative frequency of cell polarities in different E-cad and collagen concentration conditions plotted by day. (d) Percentage of cells with polarity ¿ 0.2 on days 1, 3, 5 and 7 by extracellular collagen concentration. Model predictions of the influence of extracellular collagen concentration and E-cad expression on (e) proliferation and (f) continuum invasion on days 1, 3, 5 and 7. All experimental data in (a), (b) and (d) are mean ± SEM. Statistical test used: two-way ANOVA, ****P $\leq$ 0.0001, ***P $\leq$ 0.001, **P $\leq$ 0.01, *P $\leq$ 0.05. The data in (a) and (b) are reported for N=3 for 1, 2 and 4 mg/mL collagen concentrations. The data in (c) and (d) are reported for N = 3 (1 and 2 mg/mL) or N = 2 (4 mg/mL), n = 2-4. Each technical replicate analyzed ¿100 cells. The data for 6 mg/mL are included in (a), (b), (c), and (d) as a limiting case with N=1 and n=3.

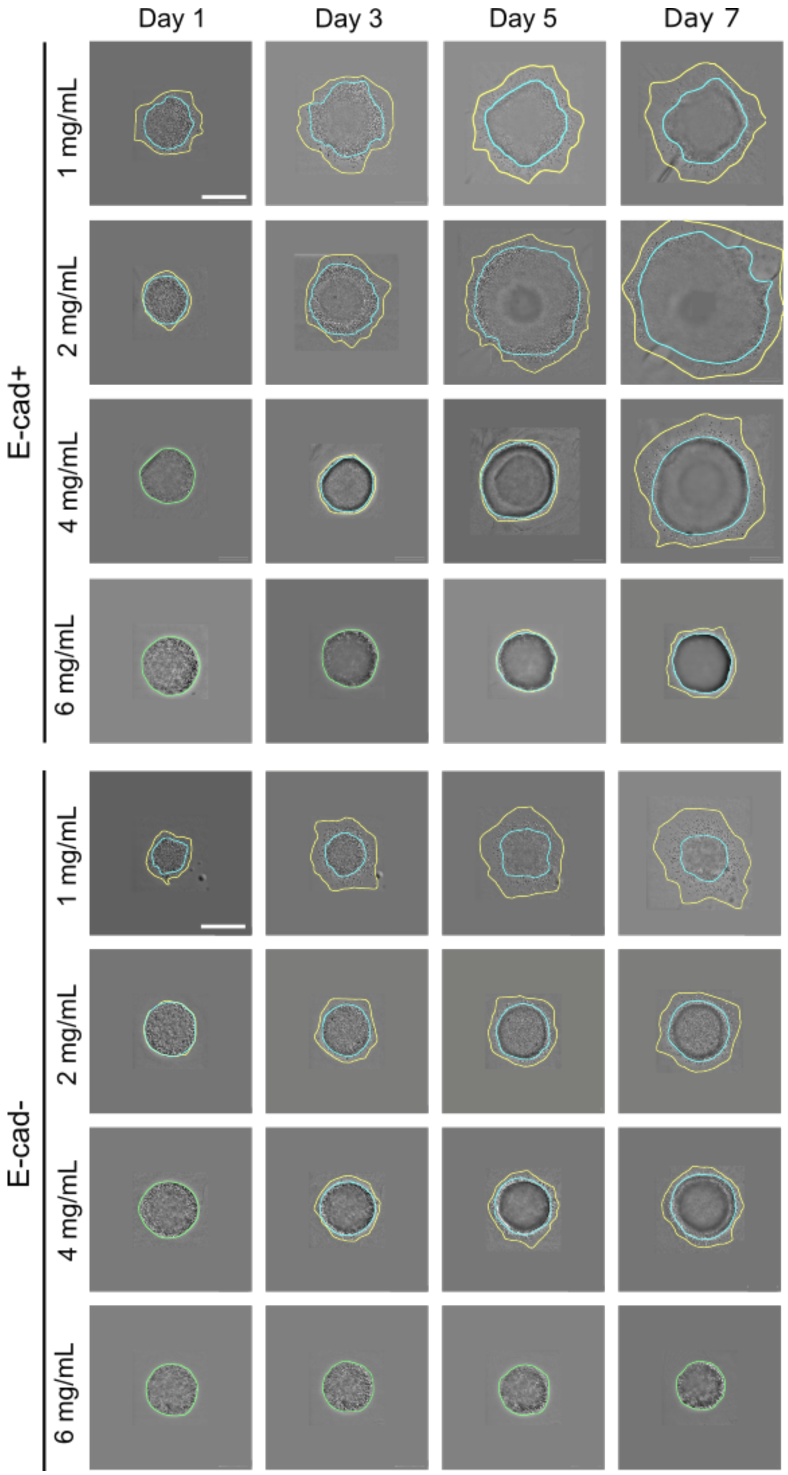


Figure S2. 2. DIC microscopy images of E-cad- and E-cad+ spheroids over time. Stitched DIC microscopy images of MDA-MB-231 cells with E-cad lentiviral knock-in (E-cad+, top), and MDA-MB-231 control cells (E-cad-, bottom) on days 1, 3, 5 and 7 with annotations of the collective region (cyan) and the detached region (yellow). Scale bars: 1 mm.

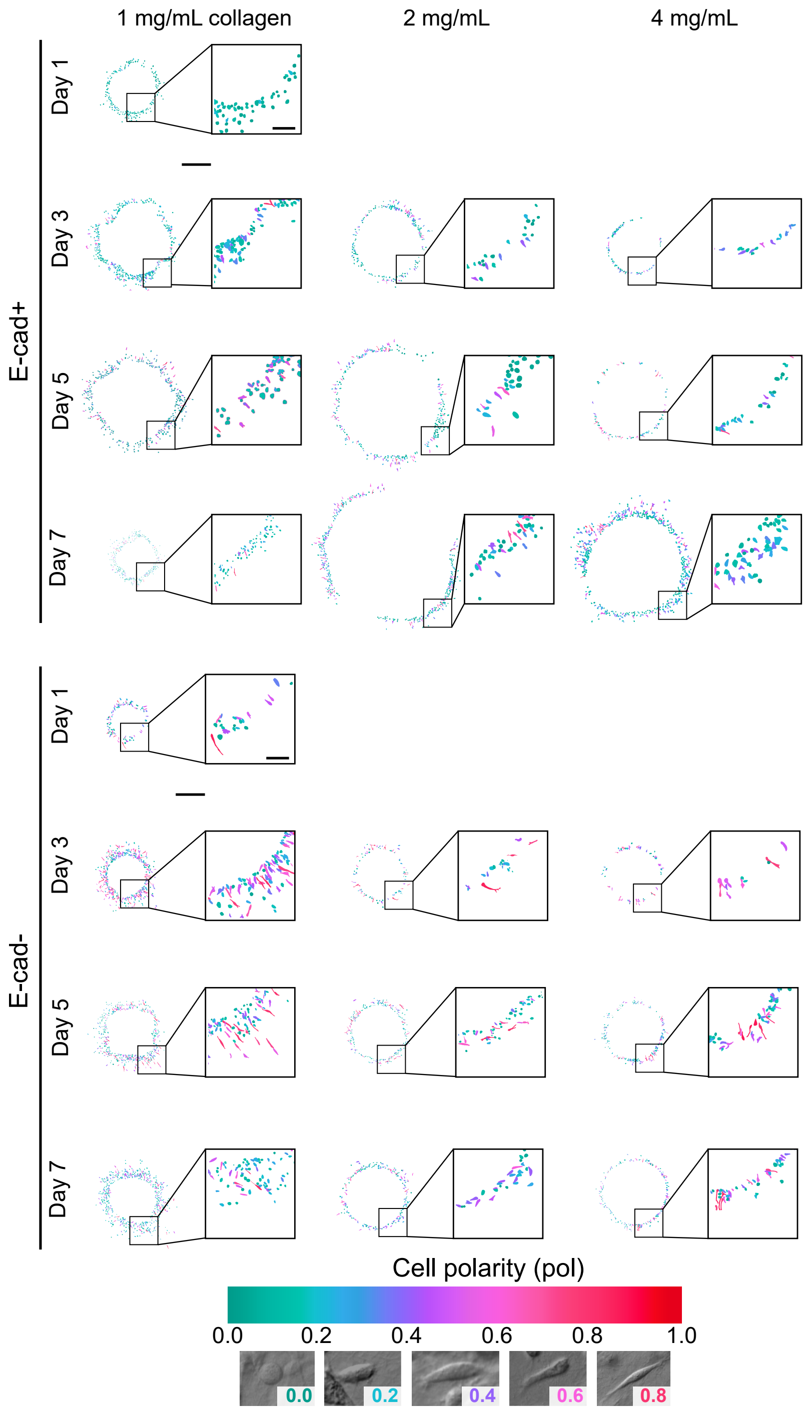


Figure S3. Cellular polarization in E-cad+ and E-cad- spheroids over time. Morphology masks of cells detached from spheroids at days 1, 3, 5 and 7 where the color represents the cell polarity score, for E-cad+ (top) and E-cad- (bottom) conditions. Examples of cells with cell polarity 0.0, 0.2, 0.4, 0.6, 0.8 and 0.9 are shown. No cells had detached from the spheroid at day 1 in the 2 and 4 mg/mL collagen matrices. N=3 biological replicates, n=2-4 technical replicates per condition. Scale bar: 1 mm, inset: 250 µm.

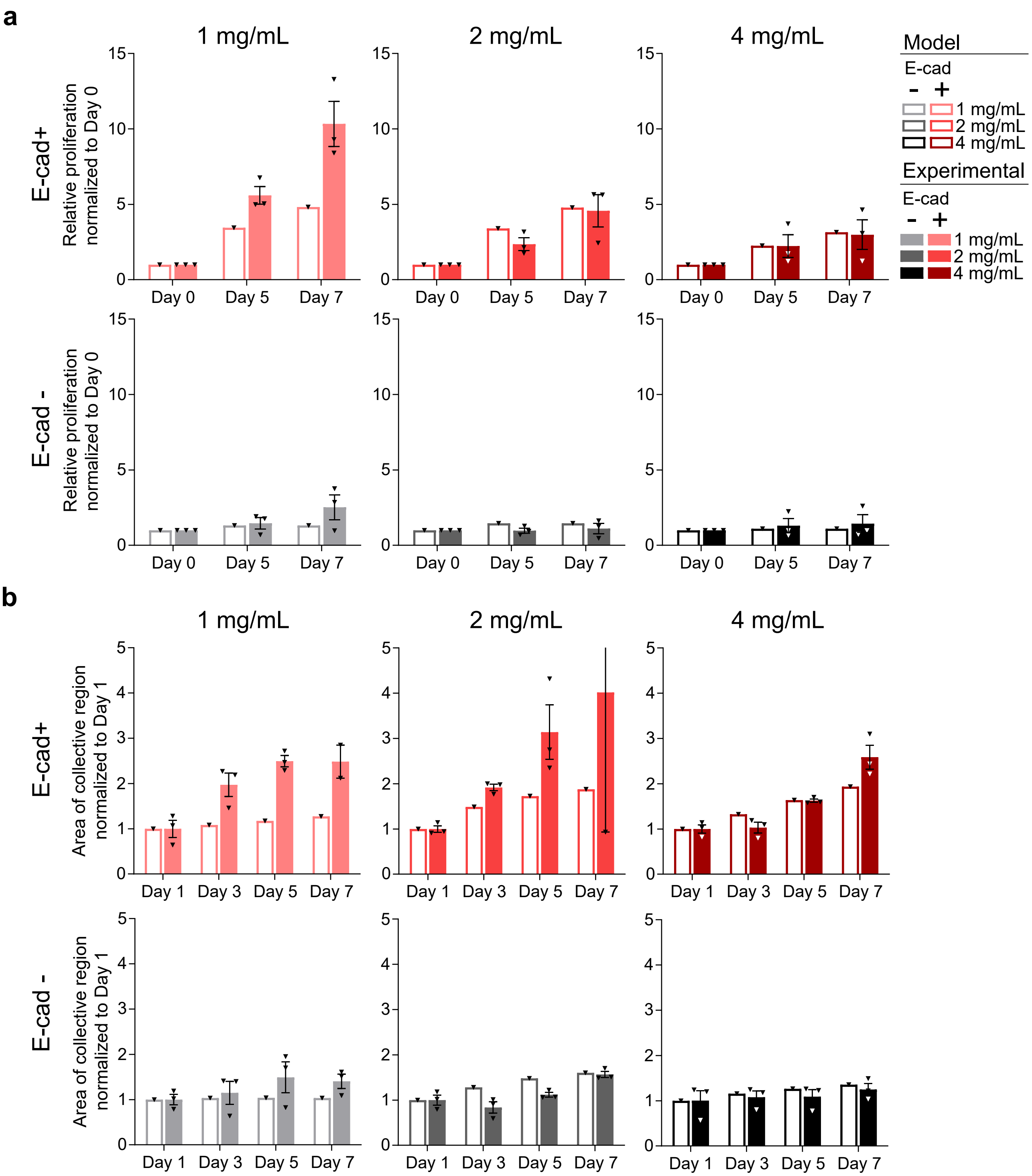


Figure S4. Comparison between experimental data and model predictions. (a) Proliferation experimental data compared to model predictions for days 0, 5 and 7. (b) Experimental data compared to model predictions of invasion dynamics for days 1, 3, 5 and 7. Error bar for experimental data on day 7 of the E-cad+, 2 mg/mL condition is cut off to improve visualization. Experimental data are mean ± SEM, N = 2-3, n = 2-4. All experimental data and model predictions are normalized to the reference case (E-cad+, 4 mg/mL collagen).

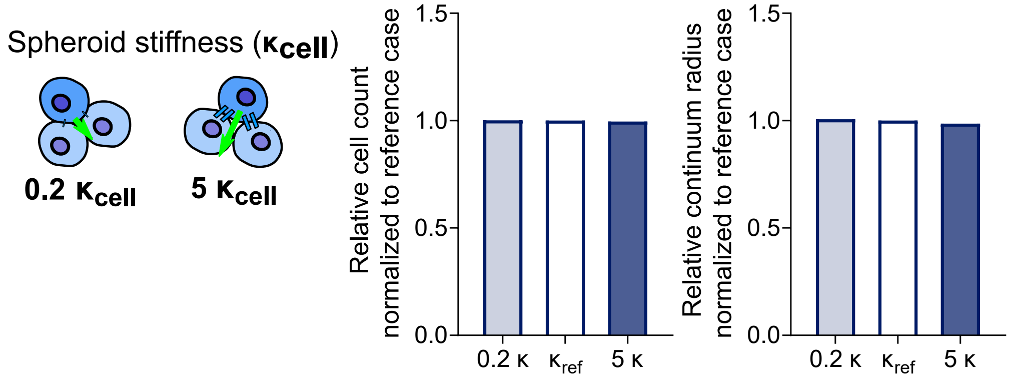


Figure S5. Parametric analysis of the model to evaluate the influence of spheroid stiffness. The reference parameters used in the simulations are those for E-cad+ and 4 mg/mL collagen concentration. The results in terms of cell number and continuum radius are normalized with the numbers obtained for the reference case. All parameters are modified dividing and multiplying the reference values by 5.

*Generalized finite element framework*

The proposed model allows for its implementation in a fully coupled finite element (FE) framework. To this end, a semi-implicit integration algorithm is formulated. Hereafter, the formulation is presented in index notation to facilitate the implementation. Indexes with capital letters, i.e., $\{I, J, K, L\}$, represent material coordinates; while indexes with lower case letters, i.e., $\{i, j, k, l\}$, represent spatial coordinates. The independent variables of the problem are the mechanical displacement field $\boldsymbol{u}$ and the cell density $\rho$.

*Strong forms*

The strong forms of the mechanical balance and the polarization flow rule residuals, Eqs. (1) and (3) in the main text, can be expressed in the deformed configuration $\Omega$ as:

$$\begin{aligned} Res^u_{ia} &= \sigma_{ij,j} - \zeta \dot{u}_i + f p_i = 0 \quad in\ \Omega \\ Res^{\rho} &= \dot{\rho} + \frac{\dot{J}}{J}\rho - d\left(\rho_{,j}\right)_{,j} - g\rho\left(1 - \frac{\rho}{\rho_\infty}\right) = 0 \quad in\ \Omega. \end{aligned} \tag{1}$$

Moreover, the deformation gradient can be written by means of the displacement field as:

$$F_{iJ} = u_{i,J} + I_{iJ}. \tag{2}$$

*Spatial and temporal discretization, and weak forms*

We estimate the spatial discretization of the finite element test functions ($\boldsymbol{\delta u^e}$ and $\delta\rho^e$) and trial functions ($\boldsymbol{u^e}$ and $\rho^e$) from the nodal values as:

$$\begin{aligned}\boldsymbol{\delta u^e} &= \sum_a N^u_a \boldsymbol{\delta u^e_a} \\ \boldsymbol{u^e} &= \sum_a N^u_a \boldsymbol{u^e_a} \\ \delta\rho^e &= \sum_a N^\rho_a \delta\rho^e_a \\ \rho^e &= \sum_a N^\rho_a \rho^e_a\end{aligned} \tag{3}$$

with $a$ referring to nodal values and $\boldsymbol{N^u}$ and $\boldsymbol{N^\rho}$ being the mechanical and proliferation shape functions, respectively.
The time derivation of a generic variable $\bullet$ is computed as $\frac{\partial \bullet}{\partial t} = \frac{\bullet - \bullet^t}{\Delta t}$, with $\bullet$ and $\bullet^t$ being the current variable and the variable from the previous time step. Next, the weak forms of the residuals are obtained by integrating within the reference (undeformed) configuration $\Omega_o$ as:

$$\begin{aligned}Res^u_{ia} &= \int_{\Omega_o} P_{iJ} N^u_{a,J} dV + \int_{\Omega_o} \zeta \frac{u_i - u^t_i}{\Delta t} N^u_a J dV - \int_{\Omega_o} f p_i N^u_a J dV = 0 \\ Res^\rho_a &= \int_{\Omega_o} \frac{\rho - \rho^t}{\Delta t} N^\rho_a J dV + \int_{\Omega_o} \frac{J - J^t}{\Delta t} \rho N^\rho_a dV + \int_{\Omega_o} d\rho_{,J} F^{-1}_{Jj} N^\rho_{a,J} F^{-1}_{Jj} J dV - \int_{\Omega_o} g\rho \left(1 - \frac{\rho}{\rho_\infty}\right) N^\rho_a J dV = 0\end{aligned} \tag{4}$$

where $J = det\left(\boldsymbol{F}\right)$.

*Semi-implicit integration algorithm*
We integrate the problem over time making use of an incremental iterative Newton-Raphson method that reduces the total residual to zero:

$$\begin{pmatrix}\boldsymbol{Res^u} \\ \boldsymbol{Res^\rho}\end{pmatrix} + \begin{pmatrix}\boldsymbol{K^{uu}} & \boldsymbol{K^{u\rho}} \\ \boldsymbol{K^{\rho u}} & \boldsymbol{K^{\rho\rho}}\end{pmatrix} \begin{pmatrix}\boldsymbol{du} \\ \boldsymbol{d\rho}\end{pmatrix} = \boldsymbol{0} \tag{5}$$

where $\boldsymbol{K^{ij}}$ are the stiffness matrices. The integration of the current problem also requires the computation of the proliferation deformation gradient, which reads as $\boldsymbol{F_p} = J_p^{1/3} \boldsymbol{I}$ with $J_p = det(\boldsymbol{F_p})$.
The stiffness matrices can be derived from the residuals as:

$$\begin{aligned}K^{uu}_{iakb} &= \frac{\partial Res^u_{ia}}{\partial u_{kb}} = \int_{\Omega_o} \frac{\partial \left(P_{iJ}\right)}{\partial F_{kL}} N^u_{a,J} N^u_{b,L} dV + \int_{\Omega_o} \frac{\partial \left(u_i J\right)}{\partial u_{kb}} \frac{\zeta}{\Delta t} N^u_a dV - \int_{\Omega_o} \frac{\partial \left(J\right)}{\partial u_{kb}} f p_i N^u_a dV \\ K^{u\rho}_{iab} &= \frac{\partial Res^u_{ia}}{\partial \rho_b} = \int_{\Omega_o} \frac{\partial \left(P_{iJ}\right)}{\partial \rho_b} N^u_{a,J} dV \\ K^{\rho\rho}_{ab} &= \frac{\partial Res^\rho_a}{\partial \rho_b} = \int_{\Omega_o} \frac{\partial (\rho J)}{\partial \rho_b} \frac{1}{\Delta t} N^\rho_a dV + \int_{\Omega_o} \frac{J - J^t}{\Delta t} N^\rho_a N^\rho_b dV + \int_{\Omega_o} d \frac{\partial \left(\rho_{,J} F^{-1}_{Jj} F^{-1}_{Jj} J\right)}{\partial \rho_b} N^\rho_{a,J} dV - \int_{\Omega_o} \frac{\partial \left(g\rho \left(1 - \frac{\rho}{\rho_\infty}\right) J\right)}{\partial \rho_b} N^\rho_a dV \\ K^{\rho u}_{akb} &= \frac{\partial Res^\rho_a}{\partial u_{kb}} = \int_{\Omega_o} \frac{\partial (J)}{\partial u_{kb}} \frac{\rho - \rho^t}{\Delta t} N^\rho_a dV + \int_{\Omega_o} \frac{\partial (J)}{\partial u_{kb}} \frac{\rho}{\Delta t} N^\rho_a dV + \int_{\Omega_o} \frac{\partial \left(F^{-1}_{Jj} F^{-1}_{Jj} J\right)}{\partial u_{kb}} d\rho N^\rho_{a,J} dV - \int_{\Omega_o} \frac{\partial (J)}{\partial u_{kb}} g\rho \left(1 - \frac{\rho}{\rho_\infty}\right) N^\rho_a dV.\end{aligned} \tag{6}$$

Note that the terms $\frac{\partial (P_{iJ})}{\partial F_{kL}}$, $\frac{\partial (P_{iJ})}{\partial \rho_b}$, $\frac{\partial (\rho J)}{\partial \rho_b}$, $\frac{\partial \left(\rho_{,J} F^{-1}_{Jj} F^{-1}_{Jj} J\right)}{\partial \rho_b}$ and $\frac{\partial \left(g\rho\left(1 - \frac{\rho}{\rho_\infty}\right) J\right)}{\partial \rho_b}$ depends on the constitutive equations chosen, while the remaining terms $\frac{\partial (u_i J)}{\partial u_{kb}}$, $\frac{\partial (J)}{\partial u_{kb}}$ and $\frac{\partial \left(F^{-1}_{Jj} F^{-1}_{Jj} J\right)}{\partial u_{kb}}$ are general.

*Reduction to pseudo 1D*

The problem primarily addressed in this work consists of an initial cell aggregate domain (see blue region in Figure S6) that is embedded into a spherical volume composed of collagen (see purple region in Figure S6). If we consider an ideal system of concentric spheres, the proliferation-migration problem can be reduced to pseudo 1D applying symmetries and taking a representative domain along a radial direction, see Figure S6.

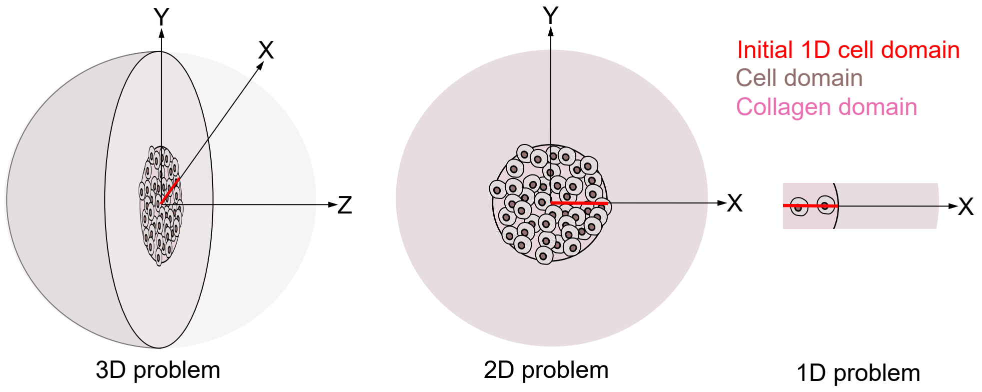


Figure S6. A schematic of the model conceptualization from 3D to pseudo 1D.

Therefore, the FE framework can be simplified to the following pseudo 1D equations:

$$Res^u = \int_{\Gamma_o} P_X N^u_{a,X} d\Gamma + \int_{\Gamma_o} \zeta \frac{u_X - u^t_X}{\Delta t} N^u_a \lambda^3 d\Gamma - \int_{\Gamma_o} f p_X N^u_a \lambda^3 d\Gamma = 0$$
$$Res^\rho = \int_{\Gamma_o} \frac{\rho - \rho^t}{\Delta t} N^\rho_a \lambda^3 d\Gamma + \int_{\Gamma_o} \frac{\lambda^3 - \lambda^3_t}{\Delta t} \rho N^\rho_a d\Gamma + \int_{\Gamma_o} d\rho_{,X} N^\rho_{a,X} \lambda d\Gamma - \int_{\Gamma_o} g\rho \left(1 - \frac{\rho}{\rho_\infty}\right) N^\rho_a \lambda^3 d\Gamma = 0 \quad (7)$$

where the subscript $X$ refers to the material coordinate along the radial direction, $\Gamma_o$ is the cell aggregate line domain and $d\Gamma = \frac{4\pi}{3}\left[X_2^3 - X_1^3\right]$ is the differential of volume in the reference configuration corresponding to a differential of radial coordinate $dL = X_2 - X_1$, with $X_i$ the local node connectivities of the element. The term $\lambda = 1 + u_{X,X}$ is the total stretch in X (radial) direction. Note that, if ideal symmetry by means of initial volumes and boundary conditions is assumed, the proliferation and migration processes develop radially leading to volumetric deformations and null shear conditions. Therefore, the effective stress, using first Piola-Kirchhoff stress, reduces to:

$$P_X = \kappa_{cell} \left(\lambda_e^3 - 1\right) \frac{\lambda_e^2}{\lambda_p} = \kappa_{cell} \left[\frac{\lambda^5}{\lambda_p^6} - \frac{\lambda^2}{\lambda_p^3}\right]. \quad (8)$$

The total radial stretch can be related to the elastic ($\lambda_e$) and proliferation ($\lambda_p$) stretches as $\lambda = \lambda_e \lambda_p$. Therefore, the stiffness matrices finally read as:

$$
\begin{aligned}
K_{ab}^{uu} &= \int_{\Gamma_o} \frac{\partial\left(P_X\right)}{\partial\lambda} N_{a,X}^{u} N_{b,X}^{u} d\Gamma + \int_{\Gamma_o} \frac{\partial\left((u_X - u_X^t)\lambda^3\right)}{\partial u_{Xb}} \frac{\zeta}{\Delta t} N_a^u d\Gamma - \int_{\Gamma_o} 3\lambda^2 f p_X N_a^u N_{b,X}^u d\Gamma \\
K_{iab}^{u\rho} &= \int_{\Gamma_o} \frac{\partial\left(P_X\right)}{\partial\rho_b} N_{a,X}^{u} d\Gamma \\
K_{ab}^{\rho\rho} &= \int_{\Gamma_o} \frac{\lambda^3}{\Delta t} N_a^\rho N_b^\rho d\Gamma + \int_{\Gamma_o} \frac{\lambda^3 - \lambda_t^3}{\Delta t} N_a^\rho N_b^\rho d\Gamma + \int_{\Gamma_o} d\lambda N_{a,X}^\rho N_{b,X}^\rho d\Gamma - \int_{\Gamma_o} g\left(1 - \frac{2\rho}{\rho_\infty}\right) N_a^\rho N_b^\rho \lambda^3 d\Gamma \\
K_{akb}^{\rho u} &= \int_{\Gamma_o} 3\lambda^2 \frac{\rho - \rho^t}{\Delta t} N_a^\rho N_{b,X}^u d\Gamma + \int_{\Gamma_o} \frac{3\lambda^2}{\Delta t} \rho N_a^\rho N_{b,X}^u d\Gamma - \int_{\Gamma_o} \lambda^2 \frac{\rho_{,X}}{\Delta t} N_a^\rho N_b^u d\Gamma \\
&+ \int_{\Gamma_o} d\rho_{,X} N_{a,X}^\rho N_{b,X}^u d\Gamma - \int_{\Gamma_o} 3\lambda^2 g\rho\left(1 - \frac{\rho}{\rho_\infty}\right) N_a^\rho N_{b,X}^u d\Gamma.
\end{aligned} \tag{9}
$$

The remaining terms to be derived read as:

$$
\begin{aligned}
\frac{\partial\left(P_X\right)}{\partial\lambda} &= \kappa_{cell}\left(\frac{5\lambda^4}{\lambda_p^6} - \frac{2\lambda}{\lambda_p^3}\right) \\
\frac{\partial\left(P_X\right)}{\partial\rho_b} &= \kappa_{cell}\left(-\frac{2\lambda^5\rho_o^2}{\rho^3} + \frac{\lambda^2\rho_o}{\rho^2}\right) N_b^\rho.
\end{aligned} \tag{10}
$$

*Model parameters used for the different simulated scenarios and additional computational results*

Table S1: Constitutive parameters used in the computational simulations.

| **Reference values for E-cad+ and 4 mg/ml collagen density** | | | | |
|---|---|---|---|---|
| $\kappa_{cell}^{compression}$ $(kPa)$ | $\kappa_{cell}^{tension}$ $(kPa)$ | $\zeta$ $(Ns/mm^4)$ | $g$ $(s^{-1})$ | $f$ $(N/mm^3)$ |
| $10^0$ | $10^{-3}$ | $2.65 \cdot 10^4$ | $1.95 \cdot 10^{-6}$ | 0.6 |

| **Friction coefficient $\zeta$ $(Ns/mm^4)$** | | | |
|---|---|---|---|
| 1 mg/ml | 2 mg/ml | 4 mg/ml | 6 mg/ml |
| $8.48 \cdot 10^2$ | $8 \cdot 10^3$ | $2.65 \cdot 10^4$ | $5.46 \cdot 10^4$ |

| **Cell polarity($\lvert\mathbf{p}\rvert$)** | | | | |
|---|---|---|---|---|
| | 1 mg/ml | 2 mg/ml | 4 mg/ml | 6 mg/ml |
| E-cad + | 0.171 | 0.166 | 0.24 | - |
| E-cad - | 0.320 | 0.412 | 0.357 | - |

| **Value adjustment for E-cad - expression** | |
|---|---|
| $\kappa_{cell}^{tension}$ $(kPa)$ | $g$ $(s^{-1})$ |
| $0.143 \cdot 10^{-3}$ | $2.65 \cdot 10^1$ |
